# Transcription factor-based gene therapy to treat glioblastoma through direct neuronal conversion

**DOI:** 10.1101/2020.08.19.257444

**Authors:** Xin Wang, Zifei Pei, Aasma Hossain, Yuting Bai, Gong Chen

## Abstract

Glioblastoma (GBM) is the most prevalent and aggressive adult primary cancer in the central nervous system (CNS). Therapeutic approaches for glioblastoma are under intense investigation, such as the emerging immunotherapy, but so far only marginal progress has been made due to the heterogeneity and highly invasive nature of glioblastoma. Here, we propose an alternative approach to tackle GBM through reprogramming proliferative GBM cells into non-proliferative neurons. We report efficient neuronal conversion from human GBM cells by overexpressing single neural transcription factor Neurogenic differentiation 1 (NeuroD1), Neurogenin-2 (Neurog2) or Achaete-scute homolog 1 (Ascl1). Subtype characterization reveals that the majority of Neurog2- and NeuroD1-converted neurons are glutamatergic, while Ascl1 favors GABAergic neuron generation. The GBM cell-converted neurons not only express pan-neuronal markers, such as NeuN and MAP2, but also exhibit neuron-specific electrophysiological activities. We further conducted transcriptome analyses to investigate the underlying cell conversion mechanism. Our RNA-seq analyses discover that neuronal genes are activated among glioma cells after overexpression of neural transcription factors, and different signaling pathways are activated by different neural transcription factors. Importantly, the neuronal conversion of GBM cells is accompanied by significant inhibition of GBM cell proliferation in both *in vitro* and *in vivo* models. Therefore, these results suggest that GBM cells can be reprogrammed into different subtypes of neurons, leading to a potential alternative approach to treat brain tumor.

**Significance:** Converting dividing glioblastoma cells into non-dividing neurons may provide an innovative therapeutic approach to treat glioblastoma.

**Highlights:**

- Efficient neuronal conversion of human glioblastoma cells achieved by overexpression of neural transcription factors
- Neurog2- and NeuroD1-converted neurons are mostly glutamatergic, while Ascl1-converted neurons are mainly GABAergic
- Transcriptome analyses reveal the activation of neuronal genes after overexpression of neural transcription factors in glioblastoma cells
- Inhibition of cell proliferation during glioblastoma cell conversion both *in vitro* and *in vivo*

## Introduction

Glioblastoma (GBM) is a type of tumor that arises from unchecked proliferation of glial cells in the brain and spinal cord ^1^. It is the most invasive primary malignant cancer in adults. In 2019, an estimated 23,820 new cases of GBM and 17,760 deaths were reported in the United States ^2^. Therapy for GBM has improved marginally over the recent years because of active proliferation, highly invasive nature, and genomic and epigenetic heterogeneity ^3,4^. Current therapeutic approaches, including immunotherapy using CART or PD-1/PD-L1 ^5–7^, try to kill or remove glioma cells entirely from the body, but this has been proved to be very difficult to achieve. Therefore, it is urgent to explore alternative therapeutic approaches for GBM.

Our lab, together with many other labs, pioneered an *in vivo* cell conversion approach to directly convert brain internal glial cells into neurons through ectopic expression of neural transcription factors ^8–12^. Considering that glioma cells originate from proliferative glial cells, we hypothesized that it might also be possible to convert glioma cells into neurons. Indeed, several studies have reported some preliminary success in such glioma cell conversion ^13–18^, but the underlying mechanisms remain largely unknown.

In this study, we tested the cell conversion capability of three different neural transcription factors, including Neurog2, NeuroD1, and Ascl1. We demonstrate that each individual factor can efficiently convert GBM cells into neuron-like cells. The converted cells not only display pan-neuronal markers but also fire action potentials, a typical electrophysiological property of neurons. Transcriptome analyses further confirmed the upregulation of neuronal genes by neural transcription factors after overexpressing in the GBM cells. Moreover, neuronal conversion of GBM cells effectively arrested cell proliferation both *in vitro* and *in vivo*. These studies suggest that transcription factor-based gene therapy may be a potential alternative approach to treat GBM.

## Results

### Efficient neuronal conversion of human glioblastoma cells

We have recently demonstrated that ectopic expression of a single neural transcription factor NeuroD1 can efficiently convert astrocytes into neurons ^9,19–25^. NeuroD1, together with Neurog2 and Ascl1, belongs to the basic helix-loop-helix (bHLH) family of neural transcription factors and play critical roles in the induction of neural differentiation during early brain development ^11,26–30^. Following our successful neuronal conversion of glial cells, we further explored the possibility of converting glioma cells into neurons using neural transcription factors. Because GBM cells are highly proliferative, we constructed retroviral vectors that can target dividing cells efficiently to overexpress Neurog2 (CAG::Neurog2-IRES-eGFP), NeuroD1 (CAG::NeuroD1-IRES-eGFP) or Ascl1 (CAG::Ascl1-IRES-eGFP) in GBM cells. Retrovirus expressing GFP alone was conducted as control (Figure 1A). After overexpressing Neurog2 or NeuroD1 in GBM cells (U251, Sigma), we found that the majority of viral infected GBM cells adopted neuronal morphology and showed immature neuronal markers such as Doublecortin (DCX) and β3-tubulin (Tuj1) at 20 days post infection (dpi), but only a small proportion of Ascl1-infected GBM cells were converted into neuron-like cells (Figure 1A). By 30 dpi, mature neuronal makers MAP2 and NeuN were both detected among Neurog2-, NeuroD1-, or Ascl1-infected cells (Figure 1B). Quantitative analyses revealed that among the 3 bHLH factors, the conversion efficiency was highest for Neurog2 (98.2% ± 0.3%), followed by NeuroD1 (88.7% ± 5.2%), and lowest for Ascl1 (24.6% ± 4.0%) at 20 dpi (Figure 1C, n = 3 repeats). At 30 dpi, the conversion efficiency was 93.2% ± 1.2% for Neruog2, 91.2% ± 1.1% for NeuroD1, and 62.1% ± 5.9% for Ascl1 (Figure 1D, n >3 repeats). Besides immunostaining approach, neuronal conversion of GBM cells was further investigated with real-time quantitative PCR (RT-qPCR) analysis to understand the time course of transcriptional changes induced by neural transcription factors. Among Neurog2-infected GBM cells, we detected a significant increase of the transcriptional activation of DCX (>2000-fold increase) at 7 dpi, which further surged to 10,000-fold change at 14-21 dpi (Figure 1E, Neurog2). NeuroD1-infected GBM cells also showed a significant increase of DCX (>1000-fold increase) at 7 dpi and over 3,000-fold increase by 14-21 dpi (Figure 1E, NeuroD1). Interestingly, Ascl1-infected GBM cells did not show significant activation of DCX until 21 dpi (Figure 1E, Ascl1), consistent with a low conversion efficiency of Ascl1 when assessed with DCX immunostaining. The low conversion efficiency by Ascl1 is not due to low expression of Ascl1 in GBM cells, because we confirmed the overexpression of Neurog2, NeuroD1 or Ascl1 with immunohistochemistry (Figure S1A), as well as RT-qPCR (Figure S1B), which actually found a huge increase of Ascl1 mRNA level among Ascl1-infected GBM cells (20 dpi). We had further performed immunostaining with immature neuronal markers DCX and Tuj1 at 6 dpi (Figure S2A-C) and found that consistent with our RT-qPCR results, Neurog2 showed highest conversion efficiency while Ascl1 showed lowest conversion efficiency. Therefore, the 3 different bHLH family neural transcription factors have different potency in converting glioma cells into neurons.

**Figure 1.**
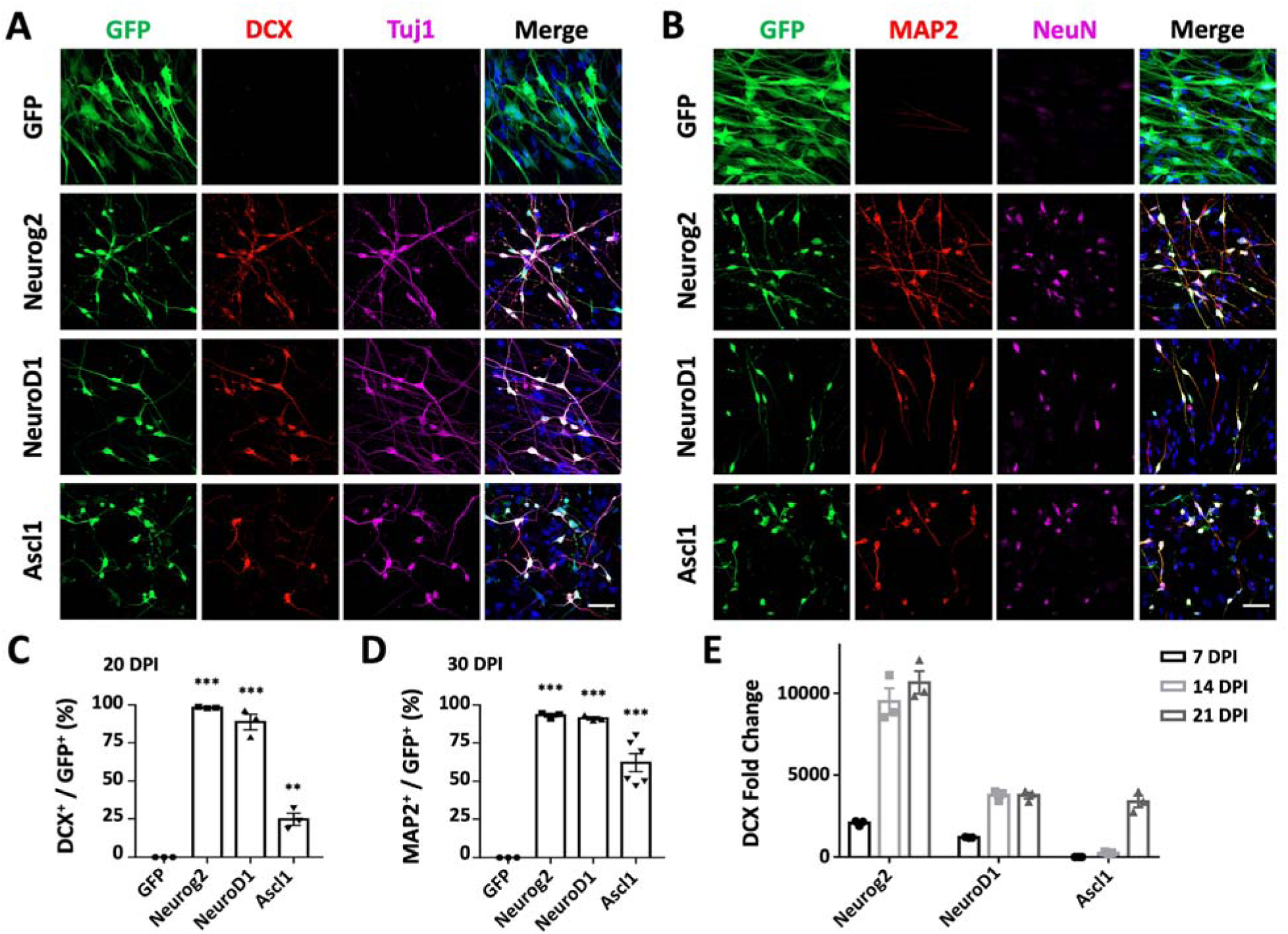
Single neural transcription factor Neurog2, NeuroD1 or Ascl1 converts human glioblastoma cells into neurons. **A-B**, Retroviral expression of Neurog2, NeuroD1 or Ascl1 in U251 human glioblastoma cells led to conversion of a large number of neuronal cells compared to the GFP control group (top row). Neurog2-, NeuroD1- or Ascl1-converted cells were immunopositive for immature neuronal markers (**A;** DCX, red; Tuj1, magenta) at 20 days post infection (dpi), and mature neuronal markers (**B;** MAP2, red; NeuN, magenta) at 30 dpi. Scale bars, 50 μm. **C-D**, Quantitative analyses of the conversion efficiency at 20 dpi (**C**) and 30 dpi (**D**). ** p < 0.01; *** p < 0.001; one-way ANOVA followed with Dunnett’s test; n ≥ 200 cells from triplicate or more cultures. **E**, Transcriptional activation of DCX during conversion revealed by real-time qPCR. Data were normalized to control GFP samples and represented as mean ± SEM. n = 3 batches.

### Characterization of the converted neurons from human glioblastoma cells

We next characterized the converted neurons from GBM cells with neuronal markers expressed in different brain regions. We found that majority of the converted cells were immunopositive for hippocampal granule neuron marker Prox1 (Figure 2A; quantified in Figure 2E: Neurog2, 90.4% ± 1.9%; NeuroD1, 89.9% ± 1.2%; Ascl1, 83.0% ± 1.4%; Prox1^+^/DCX^+^ cells), and forebrain marker FoxG1 (Figure 2B; quantified in Figure 2F: Neurog2, 99.2% ± 0.8%; NeuroD1, 87.9% ± 4.8%; Ascl1, 81.3% ± 3.6%; FoxG1^+^/MAP2^+^ cells). However, in contrast to the astrocyte-converted neurons in our previous studies^8,9,31,32^, few neurons converted from GBM cells expressed cortical neuron marker Ctip2 or Tbr1 (Figure S3A-B). These results suggest that the intrinsic imprinting of human glioblastoma cells may be different from astroglial cells and may influence the cell fate after conversion. To directly test this hypothesis, we performed side-by-side comparison with neurons converted from human astrocytes (HA1800, ScienCell, San Diego, USA). Majority of the Neurog2-, NeuroD1- or Ascl1-converted neurons from human astrocytes were positive for FoxG1 and Prox1, with a significant proportion immunopositive for Ctip2 (Figure S4A-B). Therefore, neurons converted from GBM cells shared some common properties with the neurons converted from astrocytes but differed in the specific neuronal subtypes.

**Figure 2.**
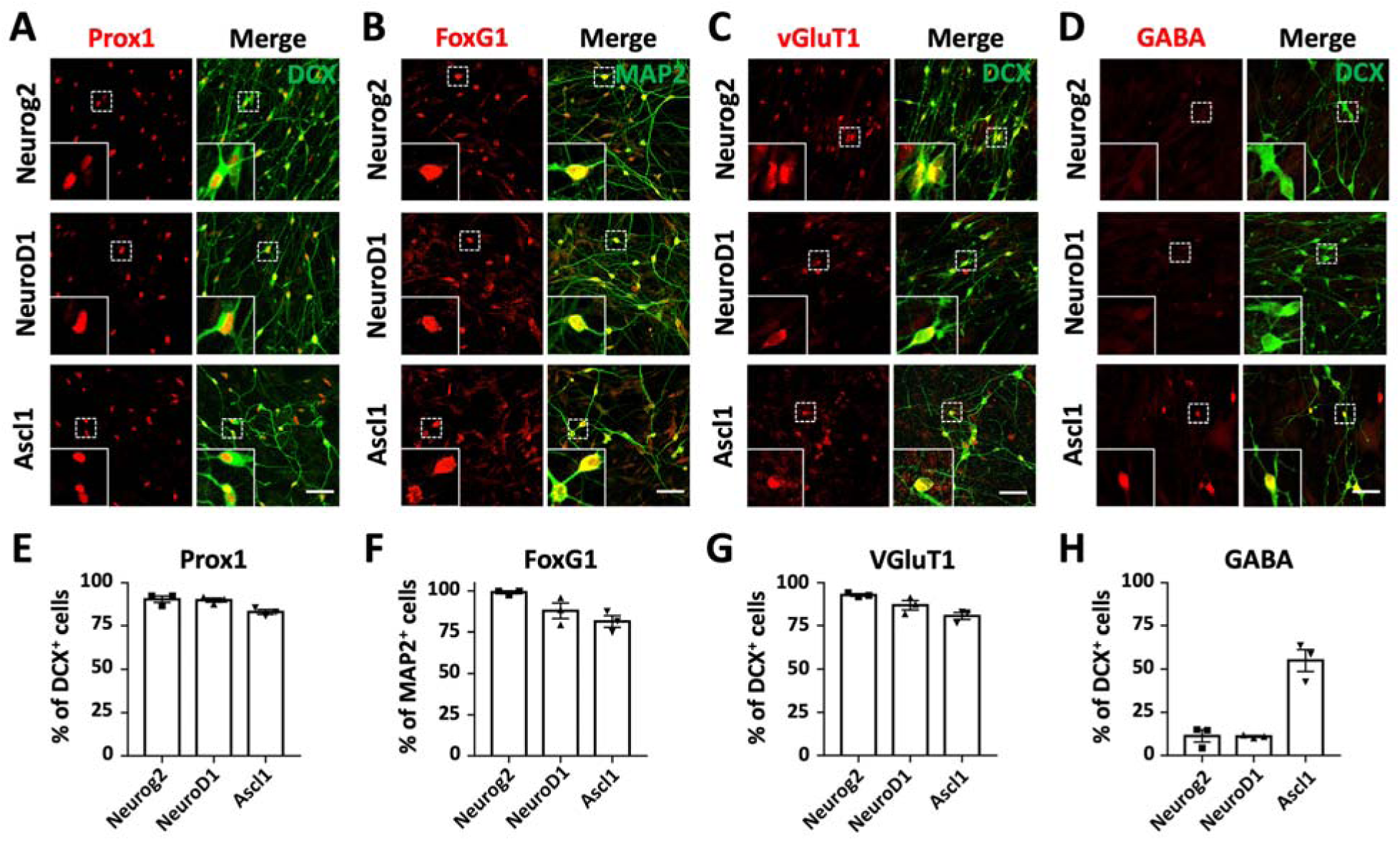
Characterization of the converted neurons from human GBM cells. **A-D**, Representative images showing the immunostaining of neuronal subtype markers in the converted neurons from U251 human GBM cells. Most of the Neurog2-, NeuroD1- and Ascl1-converted neurons (DCX or MAP2, green) were immunopositive for hippocampal neuron marker Prox1 (red in **A**), forebrain neuron marker FoxG1 (red in **B**) and Glutamatergic neuron marker VGluT1 (red in **C**). Note that there were many GABA^+^ neurons (red in **D**) converted by Ascl1 instead of Neurog2 or NeuroD1. **E-H**, Quantitative analyses of the converted neuron subtypes. Samples were at 20 dpi. Scale bars, 50 μm. Data were represented as mean ± SEM. n ≥ 200 cells from triplicate cultures.

Next, we characterized the converted neuronal subtypes according to the neurotransmitters released, in particular glutamatergic and GABAergic neurons, which are the principal excitatory and inhibitory neurons in the brain respectively. Most Neurog2-, NeuroD1-, and Ascl1-converted cells were immunopositive for glutamatergic neuron marker VGluT1 (Figure 2C; quantified in Figure 2G: Neurog2, 92.8% ± 0.7%; NeurD1, 86.9% ± 2.7%; Ascl1, 80.6% ± 2.1%; VGluT1^+^/DCX^+^ cells). The majority of Neurog2- and NeuroD1-converted cells were immunonegative for GABA (Figure 2D; quantified in Figure 2H: Neurog2, 11.1% ± 3.8%; NeuroD1, 8.6% ± 2.5%; GABA^+^/DCX^+^ cells). In contrast, roughly half of the Ascl1-converted cells were GABA-positive neurons (Figure 2D; quantified in Figure 2H: Ascl1, 55.0% ± 6.4%, GABA^+^/DCX^+^ cells), reflecting the differences among different neuronal conversion factors.

In summary, the majority of the Neurog2- or NeuroD1-converted neurons are forebrain glutamatergic neurons, while Ascl1 shows a trend for GABAergic neuron generation. Therefore, expression of different transcription factors will have significant influence on the converted neuronal subtypes.

### Fate change from glioblastoma cells to neurons induced by Neurog2 overexpression

Considering that Neurog2 yielded the fastest and most efficient neuronal conversion in GBM cells, we further investigated the Neurog2-induced conversion process in detail. Previous studies reported that the astrocyte marker GFAP and the epithelial-mesenchymal transition (EMT) marker vimentin were both highly expressed in human U251 GBM cells ^33,34^. After Neurog2 overexpression for 20 days, both GFAP and vimentin were downregulated compared to the control (Figure 3A). The gap junction marker Connexin 43 was also downregulated after Neurog2 overexpression (Figure 3B; quantified Connexin 43 intensity in Figure 3C: GFP control, 19.4 ± 0.7 a.u.; Neurog2, 11.6 ± 0.8 a.u.; at 20 days post infection), consistent with the fact that neurons have less gap junctions compared with glial cells ^35^. Interestingly, we observed typical growth cone structure among the Neurog2-converted neurons (Figure 3D), which showed fingerlike filopodia labeled by filamentous actin (F-actin) probe Phalloidin and growth cone marker GAP43 (Figure 3D).

**Figure 3.**
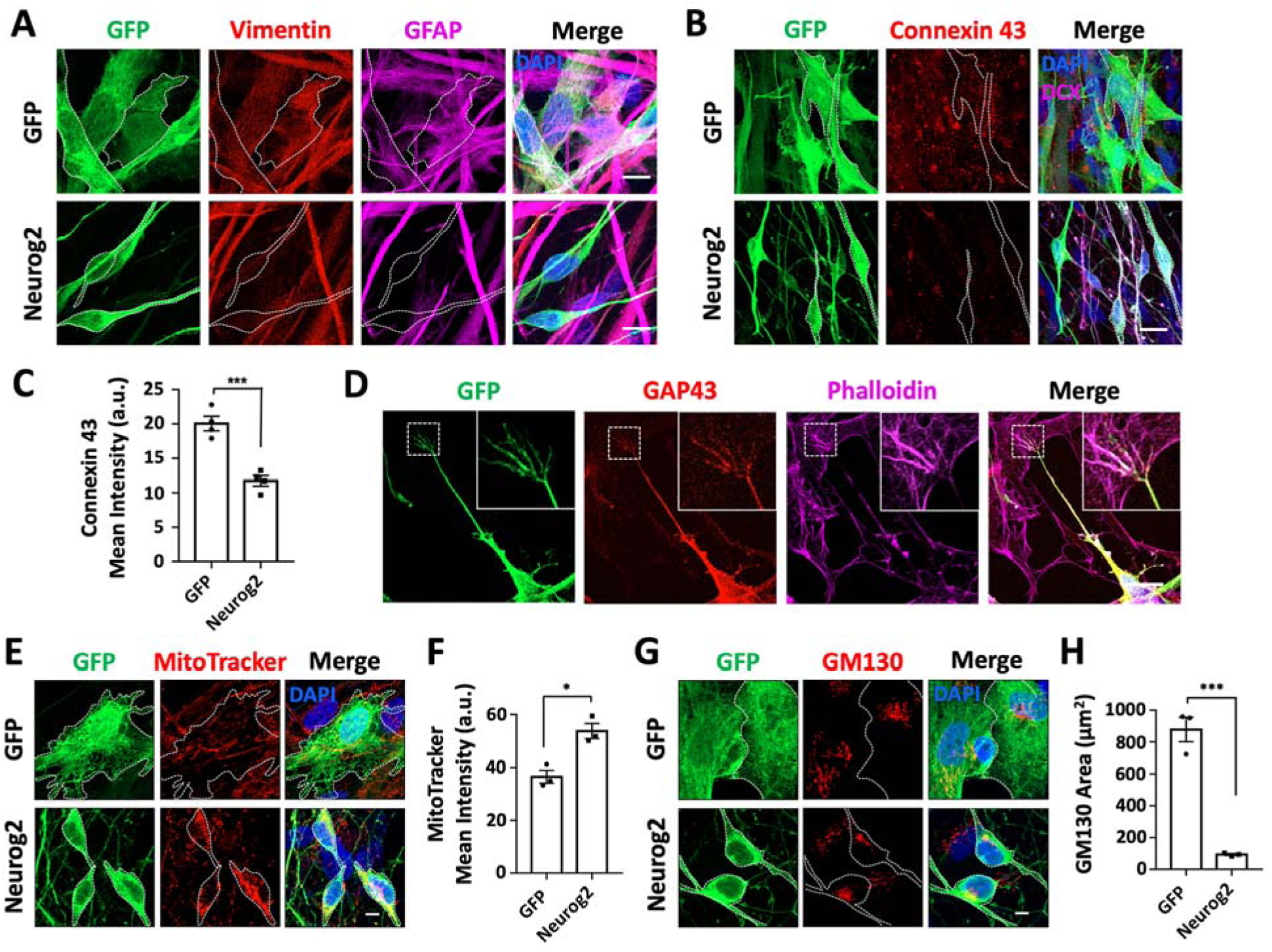
Fate change from glioblastoma cells to neurons induced by Neurog2 overexpression. **A**, Downregulation of astrocyte markers vimentin (red) and GFAP (magenta) in Neurog2-converted neurons (bottom row) compared with the control GFP-expressing U251 cells (top row). Samples were at 20 dpi. **B**, Representative images showing the gap junctions (Connexin 43, red) in U251 glioblastoma cells expressing GFP alone (top row) or Neurog2 (bottom row) at 20 dpi. **C**, Quantified data showing a significant reduction of Connexin 43 mean intensity in Neurog2-converted neurons compared with control cells. Samples were at 20 dpi. n ≥ 60 from triplicate cultures. **D**, Representative images illustrating the fingerlike filopodia of a growth cone depicted by growth cone marker Growth Associated Protein 43 (GAP43, red) and filamentous actin (F-actin) probe Phalloidin (magenta) in Neurog2-converted neurons. Samples were at 6 dpi. **E-H**, Distribution and morphological changes of mitochondria (MitoTracker, red in **E**) and the Golgi apparatus (GM130, red in **G**) during Neurog2-induced neuronal conversion in U251 cells. Quantified data showing the MitoTracker mean intensity (**F**) and the Golgi apparatus size reflected by GM130 covered area (**H**). Samples were at 30 dpi. n ≥ 150 from triplicate cultures. Scale bars, 20 μm in **(A), (B)**, and **(D)**; 10 μm in (**E**) and (**G**). Data were represented as mean ± SEM and analyzed by Student’s *t*-test. *, p < 0.05; ***, p < 0.001.

Different from GBM cells, neurons are highly polarized cells. We wondered what would happen to the cellular organelles such as mitochondria and Golgi apparatus during cell conversion given their important roles in maintaining cellular functions and homeostasis. Mitochondria are known to locate in areas with high-energy demand. In GBM cells, mitochondria distributed in the cytoplasm without obvious polarization; whereas after Neurog2-induced conversion, mitochondria showed a significant change in the distribution pattern with a concentrated localization in the soma surrounding the nucleus (Figure 3E). Compared to the control at 30 dpi, the mean intensity of mitochondria significantly increased in the converted neurons (Figure 3F). Similarly, the distribution of Golgi apparatus also showed a significant change between Neurog2-converted neurons and control GBM cells (Figure 3G). Compared to the control group, the area of Golgi apparatus was much smaller in Neurog2-converted neurons (Figure 3H). On the other hand, autophagy activity was found to be comparable between the Neurog2-converted cells and the control cells (Figure S5A-C). Together, the subcellular distribution patterns of cellular organelles undergo a significant change during the cell conversion process from GBM cells to neurons.

### Functional analyses of neurons converted from human glioblastoma cells

A critical factor for testing neuronal conversion is whether the GBM cell-converted cells can form neuronal connections and exhibit functional properties. We investigated the capability of the Neurog2-converted cells in forming synapses by performing immunostaining for synaptic vesicle marker SV2. We detected intensive synaptic puncta along MAP2-labeled dendrites in the Neurog2-converted neurons at 30 dpi (Figure 4A). Patch-clamp recordings showed significant sodium and potassium currents in the converted neurons (Figure 4B-C, 30 dpi). The majority of Neurog2-converted neurons fired single action potential (14 out of 23), with a subset of the converted neurons (8 out of 23) fired multiple action potentials (Figure 4D-E). However, no spontaneous synaptic events were recorded in the Neurog2-converted neurons at 30 dpi, suggesting that the converted neurons may still be immature or perhaps the surrounding glioma cells inhibit neuronal functions. In summary, these results indicate that human GBM cells can be reprogrammed into neuron-like cells but with partial neuronal functions when surrounded by glioma cells.

**Figure 4.**
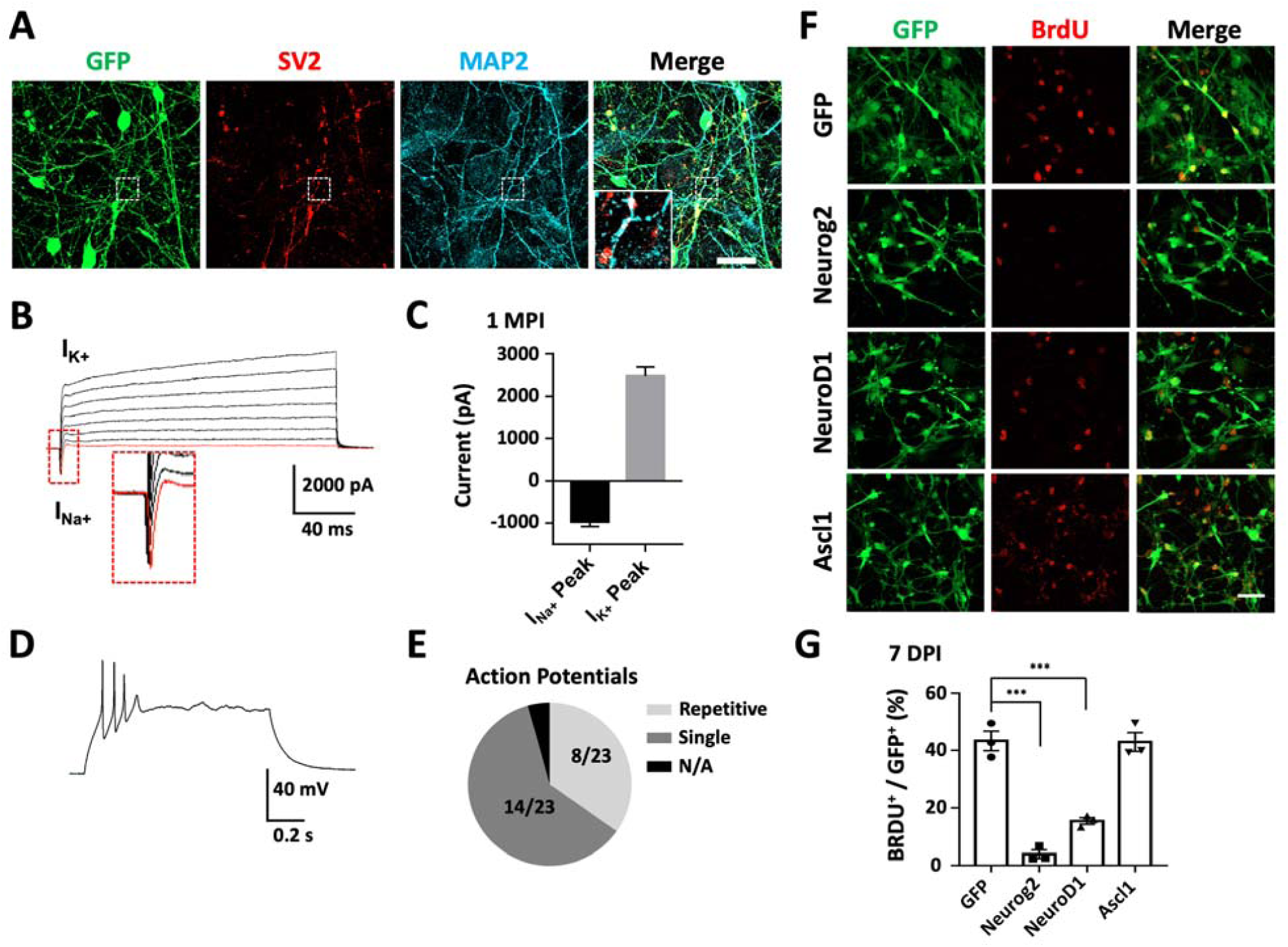
Functional analyses of human glioblastoma cell-converted neurons and cell proliferation test. **A**, Robust synaptic puncta (SV2, red) were detected along the dendrites (MAP2, cyan) in Neurog2-converted neurons from U251 human glioblastoma cells. Samples were at 30 dpi. Scale bars, 20 μm. **B-C**, Representative traces (**B**) showing Na^+^ and K^+^ currents recorded from Neurog2-converted neurons, with quantitative analyses shown in (**C**). Samples were at 30 dpi. N ≥ 20 from triplicate cultures. **D-E**, Whole-cell patch-clamp recordings revealed action potentials firing from Neurog2-converted neurons (**D**), with a pie chart indicating the fraction of cells firing single (dark grey, **E**), repetitive (light grey, **E**) or no action potential (black, **E**). Samples were at 30 dpi. n ≥ 20 from triplicate cultures. **F**, Representative images examining cell proliferation through BrdU immunostaining (red) in U251 human glioblastoma cells expressing GFP, Neurog2, NeuroD1 or Ascl1 (green). Cell cultures were incubated in 10 mM BrdU for 24 hours before immunostaining at 7 dpi. Scale bars, 50 μm. **G**, Quantitative analyses of the proliferative cells (BrdU^+^ cells/total GFP^+^ infected cells) at 7 dpi. Data were analyzed by one-way ANOVA followed with Dunnett’s test. ***, p < 0.001; n ≥ 200 cells from triplicate cultures. Data were represented as mean ± SEM.

### Arrest of cell proliferation through cell conversion

Neurons are terminally differentiated non-proliferating cells. Therefore, neuronal reprogramming may be a promising strategy to control cancer cell proliferation. To test this hypothesis, we examined cell proliferation at the early stage of conversion. GBM cells at 7 days post viral infection were incubated with 10 mM BrdU for 24 hours to label the proliferative cells (Figure 4F). Quantification of the percentage of BrdU positive cells showed that compared with GFP control, the proliferation of Neurog2- and NeuroD1-infected cells decreased significantly (Figure 4G: GFP, 64.8% ± 4.1%; Neurog2, 11.9% ± 2.9%; NeuroD1, 24.5% ± 2.4%; 7 dpi). However, the proliferation of Ascl1-converted cells remained active at 7 dpi (Figure 4F; quantified in Figure 5G: Ascl1, 54.6% ± 1.2%), possibly due to a slow action of Ascl1 in GBM cells (Figure 1A-D, S2A-C). Overall, the proliferation rate of GBM cells was significantly decreased with Neurog2 or NeuroD1 overexpression, consistent with the fast converting speed by Neurog2 and NeuroD1 after infecting GBM cells. These results suggest that in addition to neuronal conversion, ectopic expression of neuronal transcription factors may also be a promising approach to control GBM cell proliferation.

**Figure 5.**
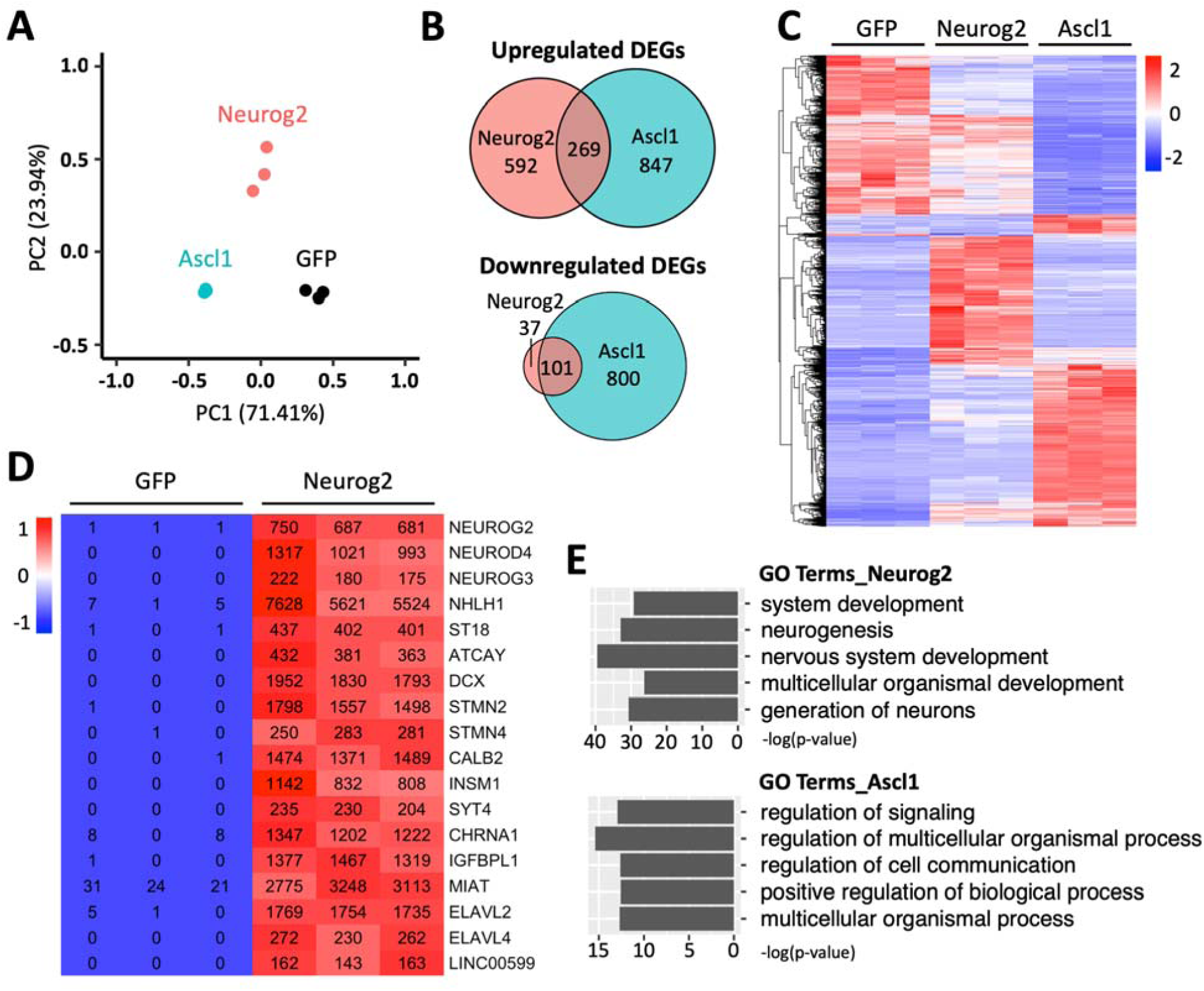
Transcriptome analyses of human glioblastoma cells with Neurog2 or Ascl1 overexpression. **A**, Principal component analysis (PCA) comparing the global gene expression profiles of U251 human glioblastoma cells infected by Neurog2, Ascl1, or control GFP retroviruses. Samples were collected at 5 DPI. n = 3 biological replicates for each group. Note that Ascl1 samples are very much clustered together. **B**, Venn diagrams showing the number of differentially expressed genes (DEGs) (adjusted p < 0.05, fold change >2, compared with control GFP samples) in Ascl1 or Neurog2 group. **C**, A heat map with hierarchical clustering showing the DEGs (adjusted p < 0.05, fold change >2, total 2,612) in response to Ascl1 or Neurog2 overexpression in U251 cells. Color scaled within each row. Normalized read count values were presented. **D**, A heat map illustrating that the top upregulated DEGs in Neurog2 samples were closely related with neurogenesis. Color scaled within each row. **E**, Gene ontology (GO) terms of the upregulated DEGs (adjusted p < 0.01, fold change >3, compared with control GFP samples) in response to Neurog2 or Ascl1 overexpression.

We have investigated biomarkers of glioblastoma with or without neuronal conversion. Interestingly, glioma markers EGFR and IL13Ra2 ^36–38^ were clearly detected after neuronal conversion at 20 days post Neurog2 infection (Figure S6A-D), suggesting that the newly converted neurons may still bear certain imprint of GBM cells, at least for the early time period after conversion.

### Transcriptome analyses of human glioblastoma cell conversion

To better understand the underlying mechanisms of the glioblastoma cell-to-neuron conversion process, we performed RNA-sequencing (RNA-seq) and transcriptome analyses of GBM cells after Neurog2 or Ascl1 overexpression, with GFP alone serving as the control group (3 replicates for each group). The RNA samples were prepared at 5 DPI to capture the early responses and potential direct targets of neural transcription factors in the early stages of conversion.

The principal component analysis (PCA) showed a clear segregation of the global gene expression profiles of different groups (Figure 5A). Pair-wise differential expression analysis showed a total of 2,612 differentially expressed genes (DEGs, fold change > 2, adjusted p < 0.05) identified in Ascl1 (2,017 DEGs) or Neurog2 (999 DEGs) group in comparison with control GFP samples (Figure 5B-C). Interestingly, while both Ascl1 and Neurog2 belong to bHLH family of neural transcription factors, only a small number of DEGs (14%, 370 out of 2,612 DEGs, Figure 5B-C) were commonly regulated by both Ascl1 and Neurog2 among infected GBM cells. We then investigated the top upregulated DEGs for potential downstream targets of the transcription factors. Most of the top upregulated DEGs by Neurog2 were closely related with neurogenesis (Figure 5D). Some were well-known neural transcription factors including NEUROG3, NEUROD4, NHLH1, ST18. Neuronal genes were also strongly upregulated by Neurog2 such as DCX and Calbindin 2 (CALB2), consistent with our immunostaining results. We also identified several interesting molecular mediators associated with microRNA and RNA regulation, such as ELAV Like RNA Binding Protein 2 (ELAVL2), ELAV Like RNA Binding Protein 4 (ELAVL4) and Long Intergenic Non-Protein Coding RNA 599 (LINC00599). In contrast, the top DEGs induced by Ascl1 were not specific to the nervous system, but rather involving developmental regulation broadly, such as Bone Gamma-Carboxyglutamate Protein (BGLAP), Calcium/Calmodulin Dependent Protein Kinase II Beta (CAMK2B), and Down Syndrome Cell Adhesion Molecule (DSCAM) (Figure S7). Consistent with these findings, most of the Gene Ontology (GO) terms enriched in the Neurog2-upregulated DEGs (adjusted p < 0.01, fold change > 3, compared with GFP samples) were neurogenesis, and nervous system development (Figure 5E). In contrast, more general GO terms were found in Ascl1-upregulated DEGs (adjusted p < 0.01, fold change > 3, compared with GFP samples), such as regulation of signaling and multicellular organismal process (Figure 5E). Together, these results imply divergent transcriptome changes in response to Neurog2 or Ascl1 overexpression in human glioblastoma cells.

Next, we investigated the signaling pathways regulated by Neurog2 versus Ascl1. Gene set enrichment analysis (GSEA) revealed that Neurog2 and Ascl1 both activated the Notch signaling pathway (Figure 6A-B). However, the leading-edge subsets of genes were quite different for these two factors (Figure 6E). For instance, Ascl1 activated the expression of the receptor encoding genes NOTCH1 and NOTCH3, while Neurog2 enhanced the expression of a different branch of the Notch signaling pathway, such as the ligand encoding gene JAG1. Moreover, Neurog2 and Ascl1 showed opposite regulation of the Hedgehog signaling pathway: Neurog2 activated while Ascl1 inhibited the Hedgehog pathway (Figure 6C-D). The heatmap of the leading-edge subsets of genes confirmed this divergence (Figure 6F). Together, the transcriptome analyses confirm the neuronal fate commitment upon early expression of Neurog2 and suggest divergent molecular mechanisms between Neurog2 and Ascl1 in converting GBM cells into neurons.

**Figure 6.**
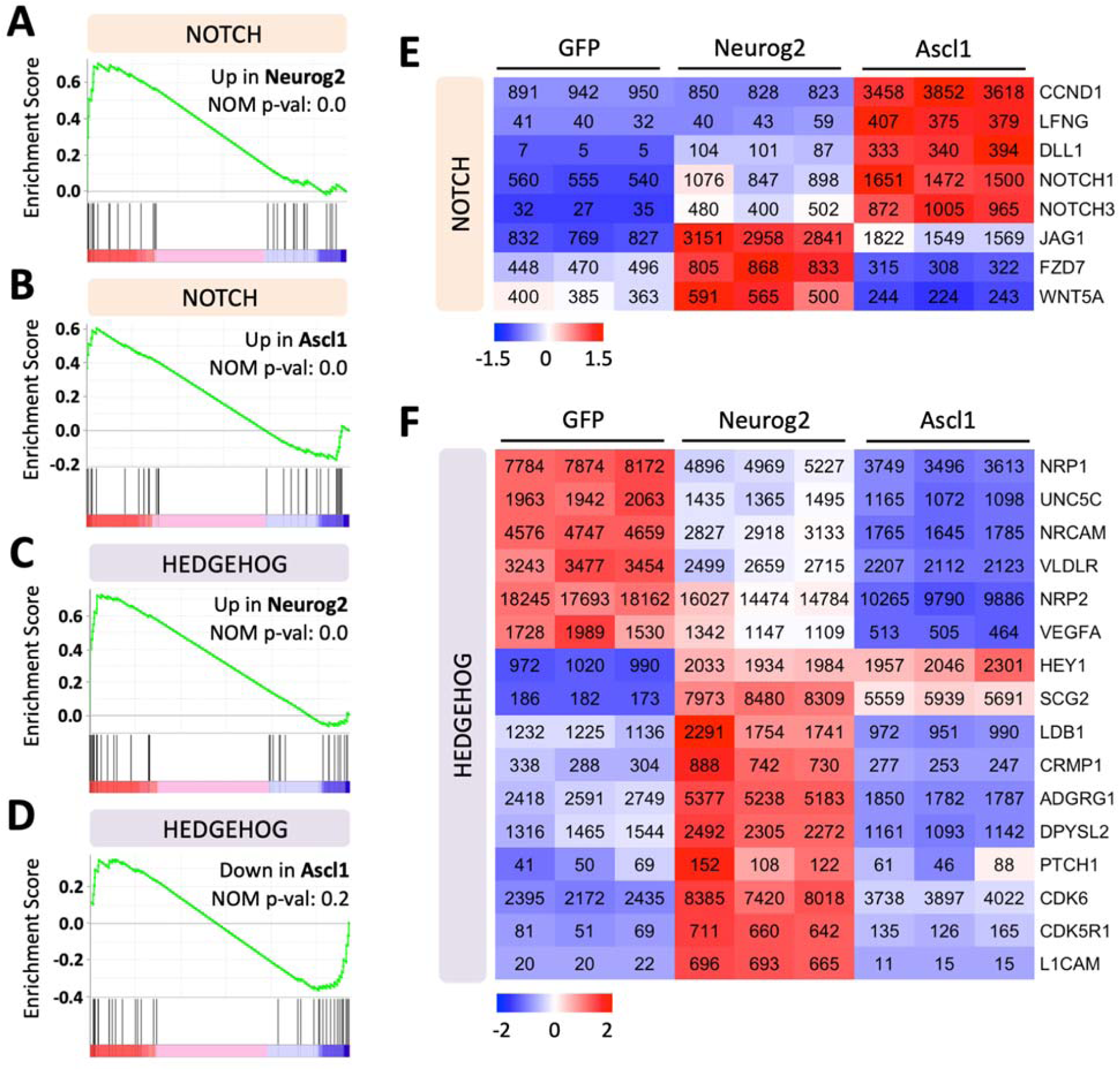
Signaling pathway changes in response to Neurog2 or Ascl1 overexpression in human glioblastoma cells. **A-D**, Gene set enrichment analysis (GSEA) of RNA-seq data showing that the Notch signaling pathway was activated in response to Neurog2 (**A**) or Ascl1 (**B**) overexpression in U251 human glioblastoma cells. In contrast, the Hedgehog signaling was significantly upregulated by Neurog2 (**C**) but not Ascl1 (**D**). **E-F**, Heat maps showing the leading-edge subsets of genes (> 100 normalized read counts at least in one sample) corresponding to the Notch (**E**) or the Hedgehog (**F**) signaling pathway shown in (**A–D**). Color scaled within each row. Normalized read count values were presented.

### *In vivo* neuronal conversion of human glioblastoma cells in a xenograft mouse model

Since *in vitro* cell culture is very different from *in vivo* environment inside the brain, we next sought to test the conversion efficiency of human glioblastoma cells in the mouse brain *in vivo*. To reduce the complication from immunorejection, we performed intracranial transplantation of human GBM cells (5×10^5^ U251 cells) into the striatum bilaterally in Rag1^-/-^ immunodeficient mice (Figure 7A). Neurog2-GFP or control GFP retroviruses with the same volume (2 μl) and titter (2 × 10^5^ pfu/ml) were injected in each side of the striatum together with the transplanted GBM cells. Transplanted GBM cells were identified by vimentin (Figure 7A) or human nuclei staining (Figure 7D). Neurog2 overexpression (Figure 7A, green cells showing the Neurog2-GFP infected GBM cells) led to an efficient neuronal conversion, indicated by immature neuronal marker DCX staining (Figure 7A-C, quantified in 7B: Neurog2, 92.8% ± 1.2%, DCX^+^/GFP^+^, 3 weeks post transplantation, n = 3 mice). Other neuronal makers such as Tuj1 and Prox1 were also detected in the Neurog2-converted neurons at one-month post transplantation (Figure 7D-E). Importantly, consistent with our *in vitro* study, the cell proliferation rate among Neurog2-infected GBM cell significantly decreased when compared to the GFP control Figure 8A-B). Unexpectedly, we observed many LCN2-positive reactive astrocytes in brain areas transplanted with GBM cells, indicating neuroinflammation after cell transplantation. However, compared to the control group, Neurog2 overexpression in the transplanted GBM cells significantly reduced the number of reactive astrocytes (Figure 8C-E), suggesting that neuronal conversion of GBM cells might ameliorate neuroinflammation in the local transplantation areas.

**Figure 7.**
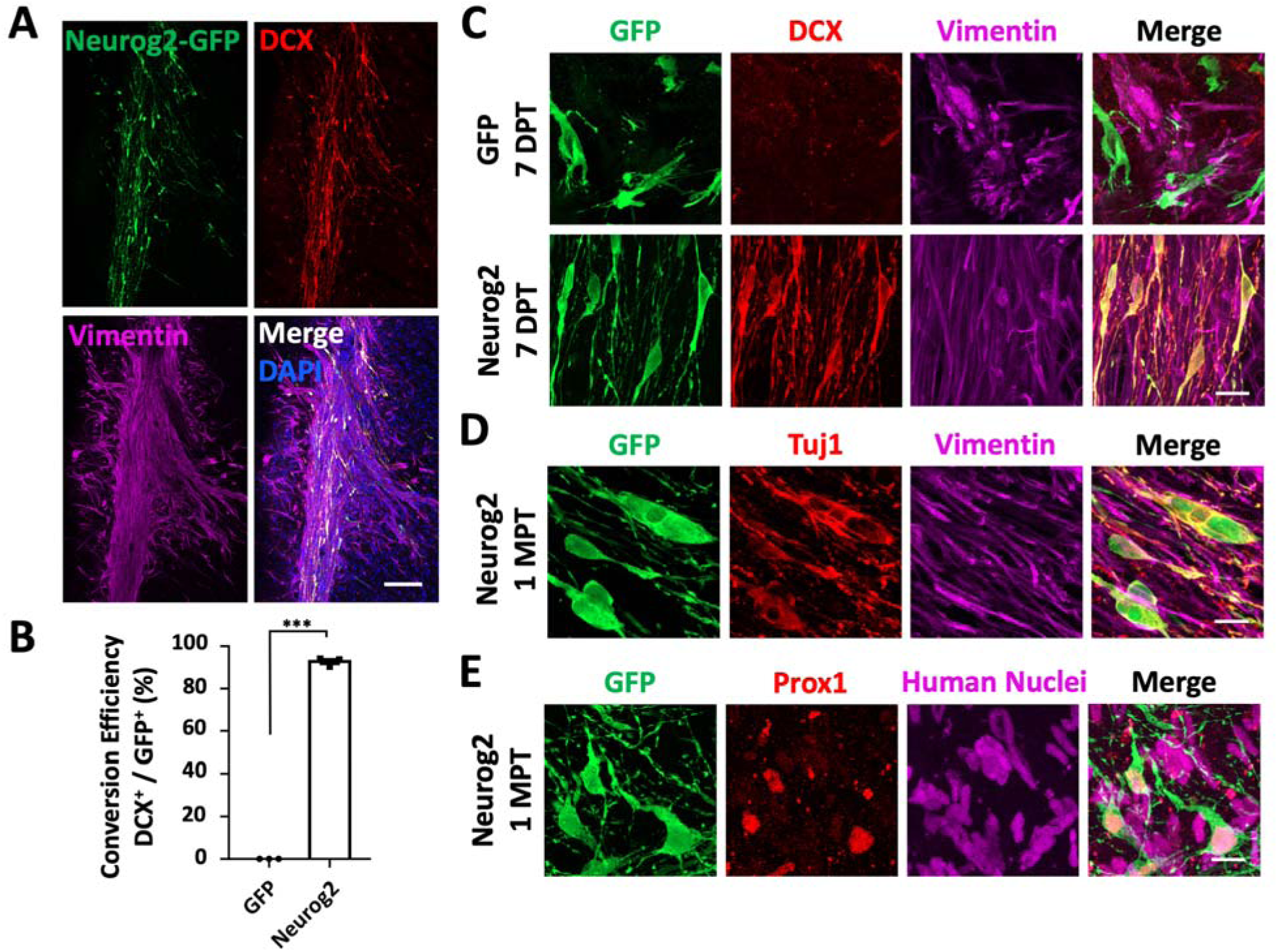
*In vivo* neuronal conversion of human glioblastoma cells in a xenograft mouse model. **A**, Representative images illustrating the transplanted human U251 GBM cells (mixed with CAG::Neurog2-IRES-eGFP retroviruses) in the striatum of Rag1^-/-^ immunodeficient mice. Note that most of the transplanted U251 cells (vimentin, magenta) transduced by Neurog2-GFP (green) retroviruses were immunopositive for neuronal marker DCX (red). Samples were at three weeks post transplantation. **B**, Quantitative analyses of the conversion efficiency at three weeks post transplantation. Data were represented as mean ± SEM and analyzed by Student’s *t*-test. ***, p < 0.001; n = 3 animals. Note that the conversion efficiency *in vivo* was also very high (92.8% ± 1.2%, DCX^+^/GFP^+^). **C**, High magnification images showing that most of the transplanted U251 cells (vimentin, magenta) infected by Neurog2-GFP (green, bottom row) retroviruses were converted into neurons (DCX, red) as early as 1-week post transplantation. **D-E**, Representative images showing that Neurog2-converted neurons (green) *in vivo* expressed neuronal markers Tuj1 (red in **D**) and Prox1 (red in **E**) at 1-month post transplantation. Transplanted U251 human glioblastoma cells were labeled by vimentin (magenta, **D**) and human nuclei (magenta, **E**). Scale bars, 200 μm in (**A**), 20 μm in (**C**)-(**E**).

**Figure 8.**
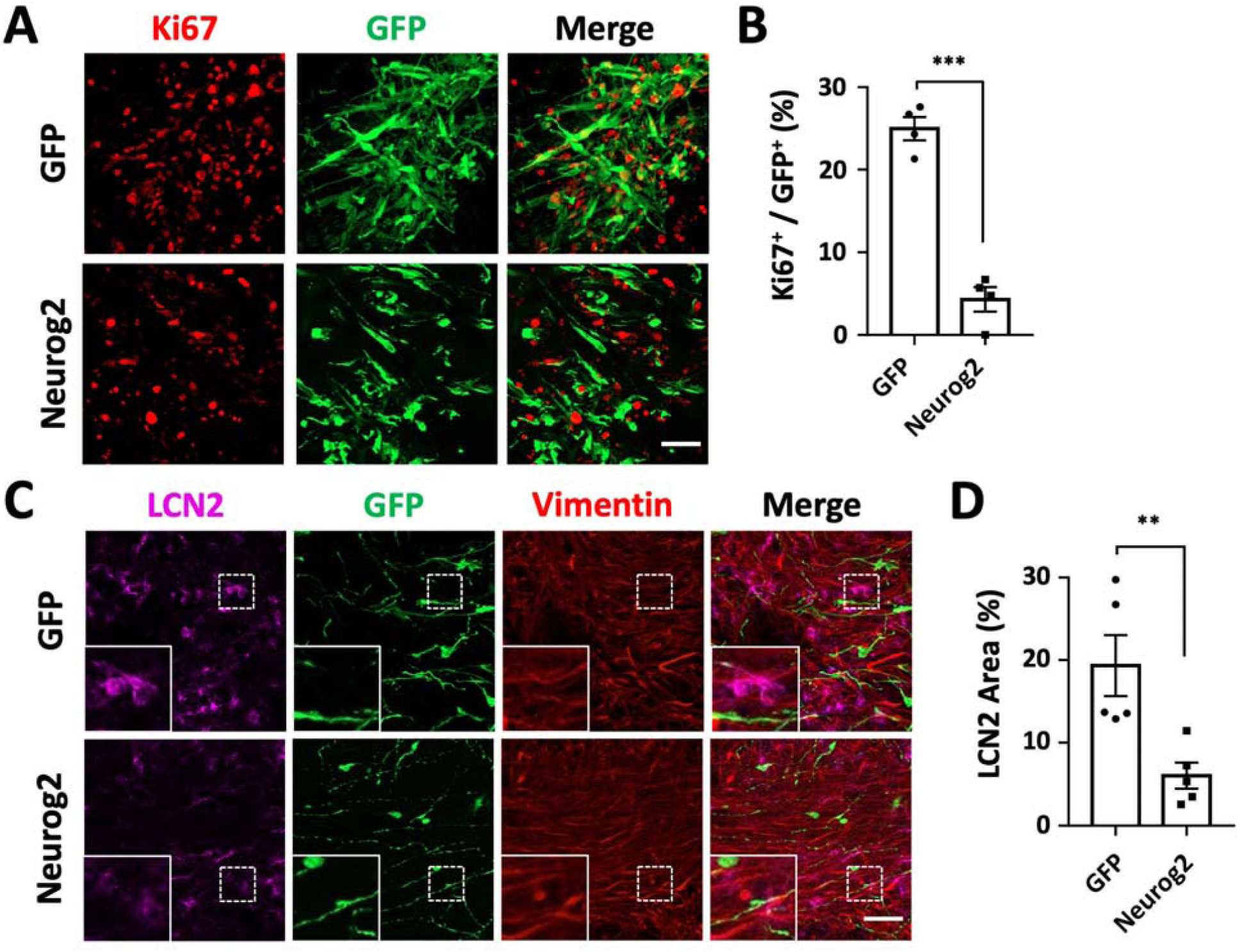
Proliferation arrest and amelioration of reactive astrocytes during *in vivo* neuronal conversion of glioblastoma cells. **A-B**, Representative images (**A**) and quantitative analyses (**B**) of proliferating U251 glioblastoma cells (Ki67^+^, red) at 7 days post transplantation. Note a significant reduction of proliferation in Neurog2-converted cells. Scale bars, 50 μm. n = 4 animals. **C-D**, Representative images (**C**) and quantitative analyses (**D**) of reactive astrocytes (labeled by LCN2, magenta) in the transplantation site with Neurog2-GFP or control GFP retroviral transduction (GFP, green). Samples were at 3 weeks post transplantation. Scale bars, 50 μm. n= 5 animals. Data were represented as mean ± SEM and analyzed by Student’s *t*-test. **, p < 0.01; ***, p < 0.001.

In summary, human glioblastoma cells can be efficiently reprogrammed into neuron-like cells through ectopic expression of neural transcription factor Neurog2 in a xenograft mouse model *in vivo*. Moreover, this reprogramming approach significantly inhibits the proliferation of glioma cells and reduces reactive astrogliosis.

## Discussion

In this study, we demonstrate that human glioblastoma cells can be converted into terminally differentiated neurons by ectopic expression of single neural transcription factor such as Neurog2, NeuroD1, or Ascl1. Remarkably, the neuronal conversion efficiency is high for all three factors tested, with Neruog2 achieving more than 90% conversion efficiency both *in vitro* and *in vivo*. More importantly, we found that during neuronal conversion of GBM cells, Neurog2 and NeuroD1 yielded more glutamatergic neurons, while Ascl1 favored GABAergic neuron generation. RNA-seq analyses confirmed the early neuronal fate commitment induced by Neurog2 and revealed divergent signaling pathways regulated by Ascl1 and Neurog2. The different neuronal subtypes induced by different neural transcription factors suggest that this cell conversion approach may have the potential to treat glioma in different brain regions enriched with different subtypes of neurons.

### Transcriptome changes in response to neural transcription factor overexpression

We conducted RNA-seq to elucidate the transcriptome changes induced by neural transcription factors in human glioma cells. Several major findings emerged from these transcriptome analyses: firstly, we discovered significant activation of neuronal genes induced by Neurog2 in human glioblastoma cells. Most of the top upregulated DEGs encode proneuronal transcription factors or well-known regulatory factors involved in neurogenesis, confirming a critical role of Neurog2 as a pioneer factor in neurogenesis. Secondly, Ascl1 and Neurog2 trigger rather distinct transcriptional changes in human glioma cells, which may explain the different neuronal fate after conversion. In fact, during embryonic development, Neurog2 and Ascl1 are involved in the generation of different neuronal subtypes in distinct regions: Neurog2 regulates the generation of glutamatergic neurons in the dorsal telencephalon, while Ascl1 is more closely related to interneuron generation in the ventral region ^39^. Consistent with their developmental functions, our transcriptome results illustrated that Ascl1 and Neurog2 elicited different neurogenic programs in human glioblastoma cells. Similar patterns were also found by other groups in neuronal conversion of human or mouse somatic cells ^29,40^. For example, Masserdotti et al. reported that after ectopic expression in astrocytes, Neurog2 activated neuronal genes involved in glutamatergic neuron maturation such as INSM1 and NeuroD4 ^29^, which were also found exclusively upregulated by Neurog2 but not Ascl1 during our human GBM cell conversion. In contrast, Ascl1 initially triggers a much broader developmental regulation program compared with the specific neuronal network activation by Neurog2. Therefore, it seems a conservative mechanism that different transcription factors will have major role in the determination of neuronal fate during neural differentiation or neural reprogramming.

### Advantages of cell conversion technology in treating glioblastoma

Glioblastoma is an aggressive cancer with highly penetrative behavior and resistant to conventional therapeutic treatment, including CART therapy ^5–7^. Traditional cancer treatment mainly aims to induce cell death, but such “cancer-killing” strategy typically produces detrimental side effects on normal cells, including epithelial cells and immune cells. The severe side effects of current chemotherapy negatively impact the quality of life for cancer patients struggling to recover from the disease and arduous treatment regimens. Converting cancer cells into non-cancerous cells is a novel therapeutic strategy that may circumvent the severe side effects of current chemotherapy or radiation therapy on normal cells ^13–18^. We demonstrate that ectopic expression of several different neuronal transcription factors can effectively convert GBM cells into non-proliferating neurons with functional properties. Moreover, the tumor cell proliferation is significantly reduced after neuronal conversion, suggesting that cancer cell conversion may be a promising strategy to control glioma. The most significant advantage of our cancer cell conversion technology is the minimal side effects on normal cells. One can envision that for small size glioma identified by MRI or other brain imaging techniques, viruses expressing neuronal transcription factors such as Neurog2 or NeuroD1 can be injected directly into the tumor to induce neuronal conversion and inhibit glioma cell proliferation. For larger size glioma, a surgical resection may be necessary, followed by injection of viruses to convert the remaining glioma cells into neurons. This approach would enable the neurosurgeons to preserve as much healthy brain tissue as possible to minimize collateral damage to the brain. More importantly, such intra-glioma injection of viral particles would have minimal side effects on normal cells such as epithelial cells or immune cells, allowing recovering patients to live a relatively normal life.

### Challenges of the cell conversion technology in cancer treatment

Like any new technology, there are certainly many challenges facing the cancer cell conversion approach. While our data suggests that cancer cell conversion therapy has the potential to slow or eradicate tumor growth, one obvious concern is that, although we have achieved high conversion efficiency (over 90% in this study), the remaining cancer cells may still pose the threat of relapse. Nevertheless, we believe such cancer cell conversion technology can significantly delay the cancer development and extend the life span of cancer patients. Also of concern is that, while retroviruses may target rapidly proliferating cancer cells, the immunoreactivity of retroviruses is problematic when considering clinical trials ^41–43^. The use of adeno-associated virus (AAV), which is a less immunogenic viral vector ^44^, needs to be further investigated as an effective delivery system to introduce neuronal transcription factors into rapidly dividing GBM cells. A third challenge would be to develop an effective system specifically targeting cancer cells for the expression of neuronal transcription factors necessary for cell conversion. Obviously, much work is needed to further address these challenges.

## Conclusion

In summary, our study showed an efficient neuronal conversion of U251 human GBM cells by forced overexpression of single neuronal transcription factor Neurog2, NeuroD1 or Ascl1. In addition, we found that this conversion approach had significant potential in generating specific types of neurons using different factors. For example, Neurog2 and NeuroD1 yielded more glutamatergic neurons, while Ascl1 favored GABAergic neuron generation. More importantly, the neuronal conversion approach resulted in a significant proliferation arrest in both cultured glioblastoma cells and a glioma xenograft mouse model. These synergistic effects of neuronal conversion plus proliferation arrest suggest a potential new therapeutic strategy to treat brain cancer. While much more studies are necessary to perfect this new technology, we envision that this unique approach of neuronal reprogramming may significantly benefit cancer patients in the future.

## Acknowledgments

We would like to thank all Chen lab members for their discussion and suggestions for this project. We thank Joseph Gyekis and Donna Sosnoski for proofreading the manuscript. This work was supported by Charles H. “Skip” Smith Endowment Fund and Verne M. Willaman Endowment Fund from the Pennsylvania State University to G.C.

## Author Contributions

G.C. conceived and supervised the entire project, analyzed the data and revised the manuscript. X.W. performed the experiments, analyzed the data, and wrote the manuscript. A.H. performed the double-blind quantifications. Z.F.P. and Y.T.B. participated in the discussion and experimental designing.

## Materials and Methods

### Cell Culture

Human GBM cell lines were purchased from Sigma (U251) or ATCC (U118). U251 cells were cultured in GBM culture medium, which included MEM (GIBCO), 0.2% penicillin/streptomycin (GIBCO), 10% FBS (GIBCO), 1mM Sodium Pyruvate (GIBCO), 1% Non Essential Amino Acids (NEAA, GIBCO), and 1 x GlutMAX (GIBCO). U118 cells were cultured in culture medium including DMEM (GIBCO), 10% FBS and 1% penicillin/streptomycin.

Human astrocytes were purchased from ScienCell (HA1800, San Diego, USA). Human astrocytes were cultured in human astrocyte medium, which included DMEM/F12 (GIBCO), 10% FBS, 3.5 mM Glucose (Sigma), and 0.2% penicillin/streptomycin, supplemented with B27 (GIBCO), N2 (GIBCO), 10 ng/ml fibroblast growth factor 2 (FGF2, Invitogen), and 10 ng/ml epidermal growth factor (EFG, Invitrogen).

For subculture, cells were trypsinized by 0.25% Trypsin (GIBCO) or TrypLE Select (Invitrogen), centrifuged for 5 min at 800 rpm, re-suspended and plated in corresponding culture medium with a split ratio around 1:4. Cells were maintained at 37°C in humidified air with 5% CO_2_.

### Reprogramming human GBM cells into neurons

U251 cells were seeded in poly-D-lysine-coated coverslips in 24-well plates at least twelve hours before the virus infection with a density of 10,000 cells per coverslip. GFP, Neurog2, NeuroD1 or Ascl1 retrovirus was added in GBM cells together with 8 μg/ml Polybrene (Santa Cruz Biotechnology). Culture medium was completely replaced by neuronal differentiation medium (NDM) the next day to help with neuronal differentiation and maturation. NDM included DMEM/F12 (GIBCO), 0.4% B27 supplement (GIBCO), 0.8% N2 supplement (GIBCO), 0.2% penicillin/streptomycin, 0.5% FBS, Vitamin C (5 µg/ml, Selleck Chemicals), Y27632 (1 µM, Tocris), GDNF (10 ng/ml, Invitrogen), BDNF (10 ng/ml, Invitrogen) and NT3 (10 ng/ml, Invitrogen). Cells were maintained at 37°C in humidified air with 5% CO_2_.

### *In vivo* neuronal conversion of human glioblastoma cells

*In vivo* neuronal conversion of human glioblastoma cells was conducted using Rag1 KO immunodeficient mice (B6.129S7-Rag1t^m1Mom^/J, The Jackson Laboratory, Stock #002216). Half a million (5×10^5^) U251 human glioblastoma cells were transplanted into the striatum of Rag1 KO mouse brains using a stereotaxic device (Hamilton). Retroviruses expressing Neurog2-GFP or GFP alone with similar titer were injected intracranially at the same time and location. Mouse brains were harvested and sliced at 1, 2, 4, and 8 weeks post injection. Immunostaining for brain slice sections were the same as cultured cells. The experimental protocols were approved by The Pennsylvania State University IACUC (IACUC # 47890).

### Next generation sequencing and data analysis

RNA was extracted using the NucleoSpin® RNA kit (Macherey-Nagel) following the manufacturer’s protocols. RNA samples came from three batches of U251 glioblastoma cells overexpressing GFP alone or Neurog2-GFP, Ascl1-GFP, making total nine samples. Quality check of RNA samples, mRNA enrichment, library construction, and next generation sequencing (single-end, 50 bp, Hiseq 3000 platform) were performed in UCLA Technology Center for Genomics and Bioinformatics. The raw data (fastq files) was checked using FastQC (v. 0.11.3) with default settings ^45^. The read alignment (against hg38 human reference genome) was performed using BWA-MEM (v. 0.7.17-r1188) and summarized using featureCounts (v. 1.5.0) ^46,47^. Differential expression analysis was processed using DESeq2 (v. 1.16.1) ^48^. DEGs in the Neurog2 or Ascl1 overexpression group were defined with adjusted p-value < 0.05, fold change > 2, compared with GFP group via DESeq2. Heatmaps were created using R console as described ^49^. Gene ontology was analyzed in Gene Ontology Consortium (http://geneontology.org/). Gene enrichment analysis was conducted using GSEA software ^50^.

Other methods can be found in the Supplemental Materials.

