## Supplemental materials for "Transcription factor-based gene therapy to treat glioblastoma through direct neuronal conversion"

Xin Wang^1,*^, Zifei Pei^1^, Aasma Hossain^1^, Yuting Bai^1^, Gong Chen^1,2,*^

**Supplemental Figures**

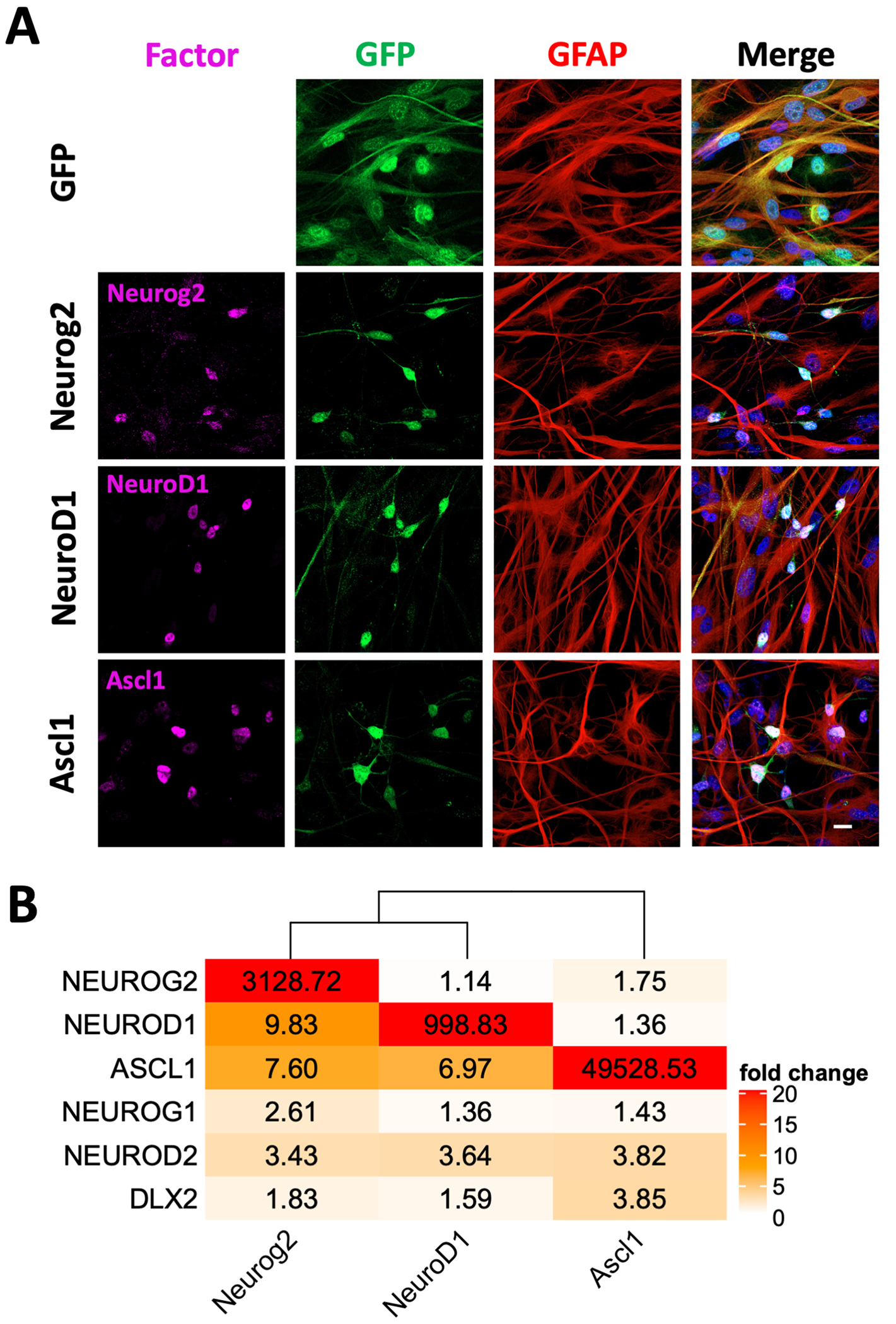

**Figure S1. Overexpression of neural transcription factor Neurog2, NeuroD1 or Ascl1 in human glioblastoma cells**

**A,** Representative images showing the immunostaining of Neurog2, NeuroD1, or Ascl1 in U251 human glioblastoma cells after retroviral transduction at 20 dpi. Scale bars, 20 μm.

**B,** A heat map summarizing the expression of different neural transcription factors revealed by real-time qPCR. Note a huge transcriptional increase of NEUROG2, NEUROD1 or ASCL1 in the corresponding group. Data were normalized to control GFP-expressing U251 cells. Data were represented as mean values from n=3 batches of cultures. Samples were collected at 20 dpi.

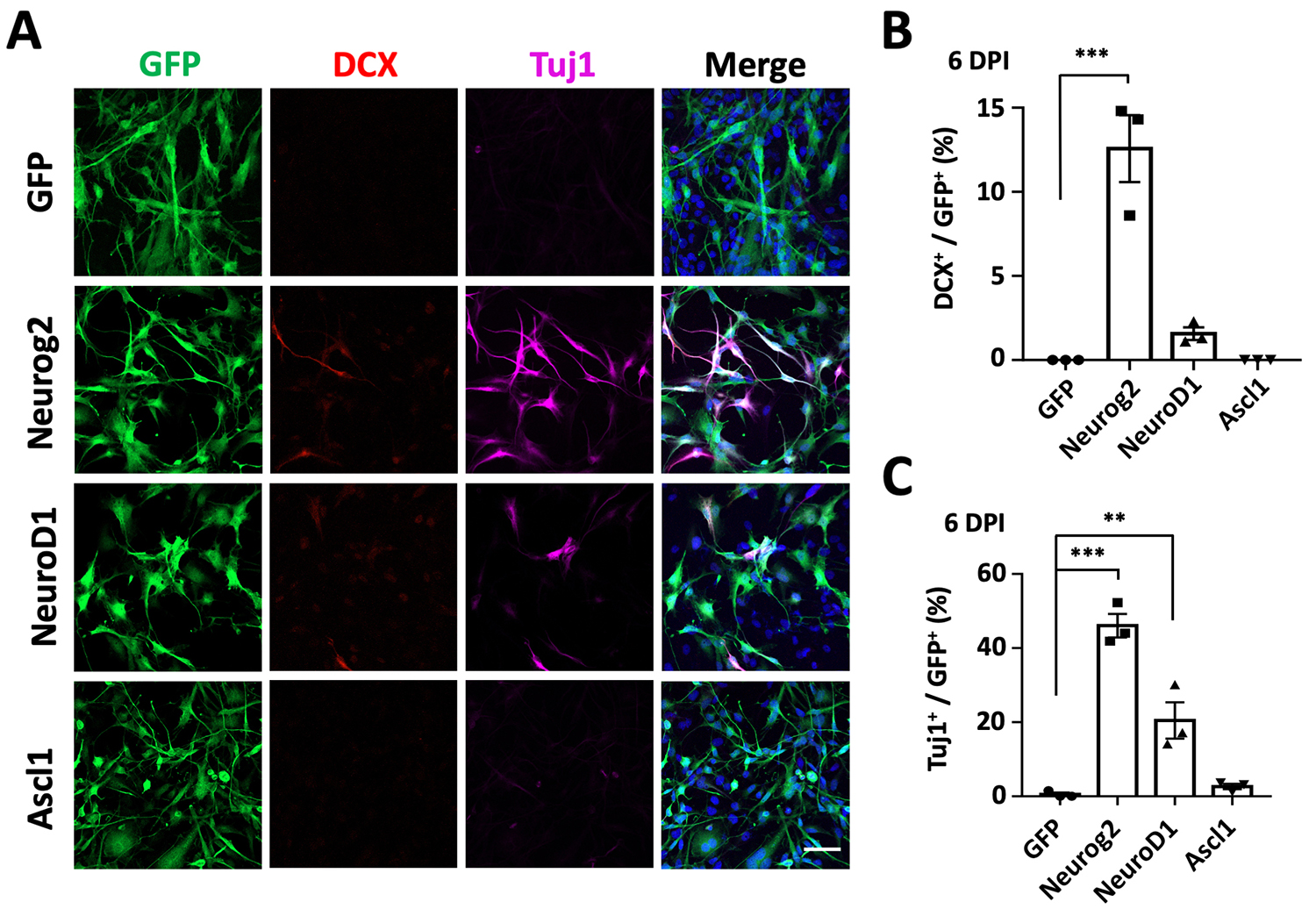

**Figure S2. Rapid neuronal conversion of human glioblastoma cells by Neurog2 and NeuroD1**

**A,** Representative images showing the immunostaining of immature neuronal markers Doublecortin (DCX, red) and β3-tubulin (Tuj1, magenta) in U251 human GBM cells with retroviral expression of Neurog2, NeuroD1, Ascl1 or GFP alone at 6 dpi. Scale bars, 50 μm.

**B-C,** Quantitative analyses of the conversion efficiency at 6 dpi. Note that Neurog2 and NeuroD1 overexpression induced a fast production of DCX^+^ cells (**B**) and Tuj1^+^ cells (**C**). Data were represented as mean ± SEM and analyzed by one-way ANOVA followed with Dunnett’s test. **, p < 0.01; ***, p < 0.001; n > 200 cells from triplicate cultures.

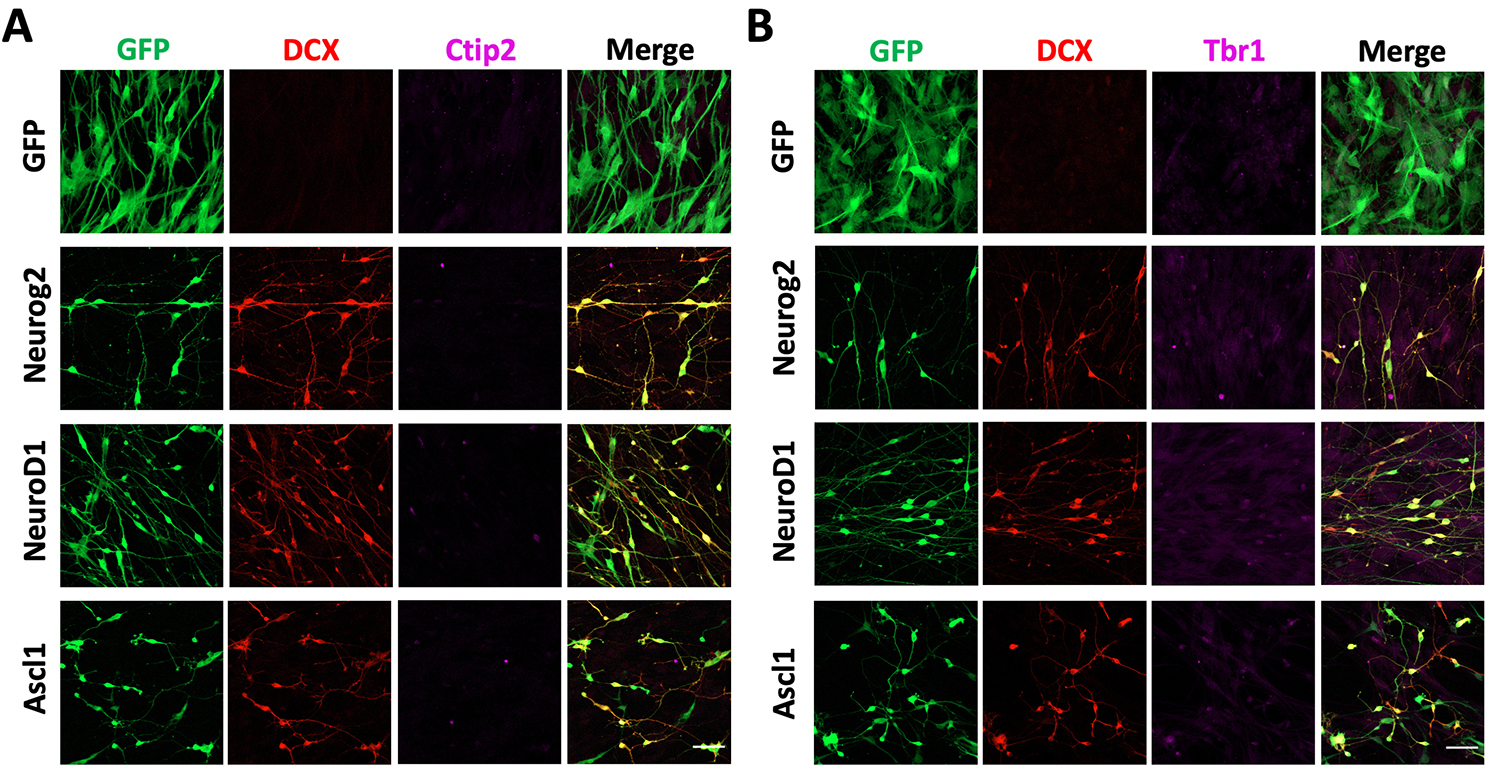

**Figure S3. Characterization of the converted neurons from human GBM cells**

**A-B,** Representative images showing the immunostaining of cortical neuron marker Ctip2 (magenta, **A**) or Tbr1 (magenta, **B**) in U251 human GBM cells overexpressing Neurog2, NeuroD1, Ascl1 or GFP alone. Samples were collected at 20 dpi. Scale bars, 50 μm.

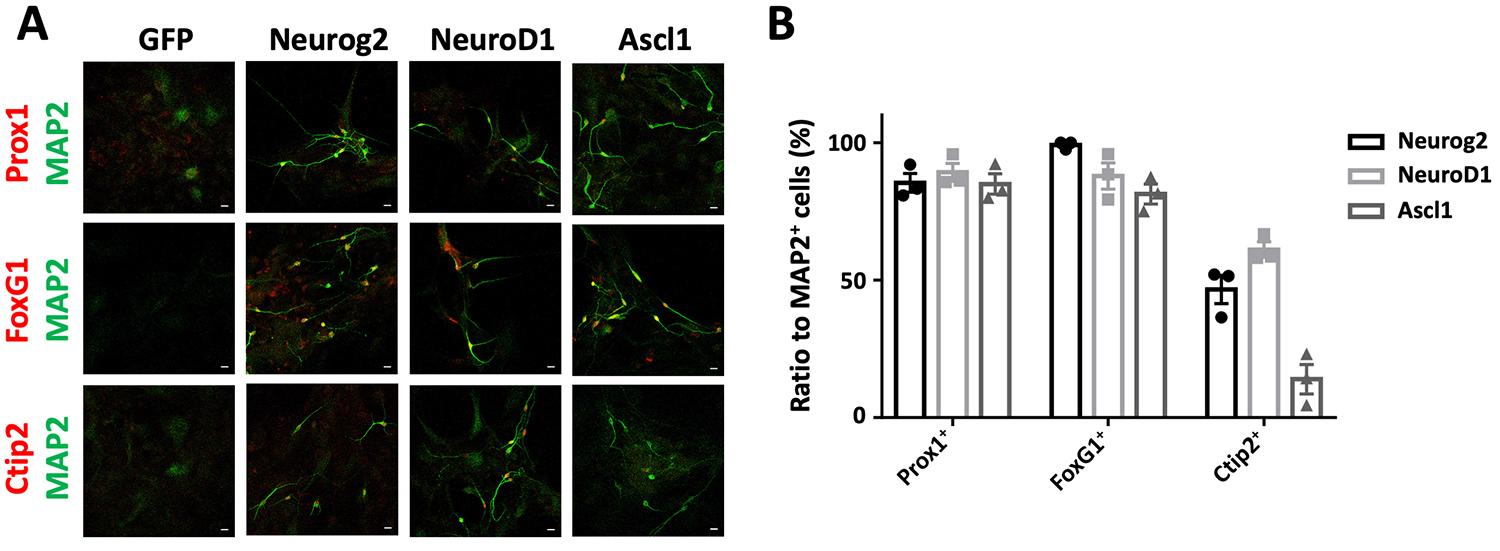

**Figure S4. Characterization of the converted neurons from human astrocytes**

**A,** Representative images showing the expression of neuronal subtype markers in the converted neurons from human astrocytes (HA1800 cells, ScienCell, San Diego, USA).

Most of the Neurog2-, NeuroD1- and Ascl1-converted neurons (MAP2, green) were immunopositive for hippocampal neuron marker Prox1 (red, top row) and forebrain marker FoxG1 (red, second row). Note that there were a decent number of Ctip2^+^ neurons (red, third row) converted by Neurog2 or NeuroD1. Scale bars, 20 μm.

**B**, Quantitative analyses of Neurog2-, NeuroD1- and Ascl1-converted neurons from human cortical astrocytes (HA1800 cells, ScienCell, San Diego, USA).

Samples were at 30 dpi. Data were represented as mean ± SEM. n > 50 cells from triplicate cultures.

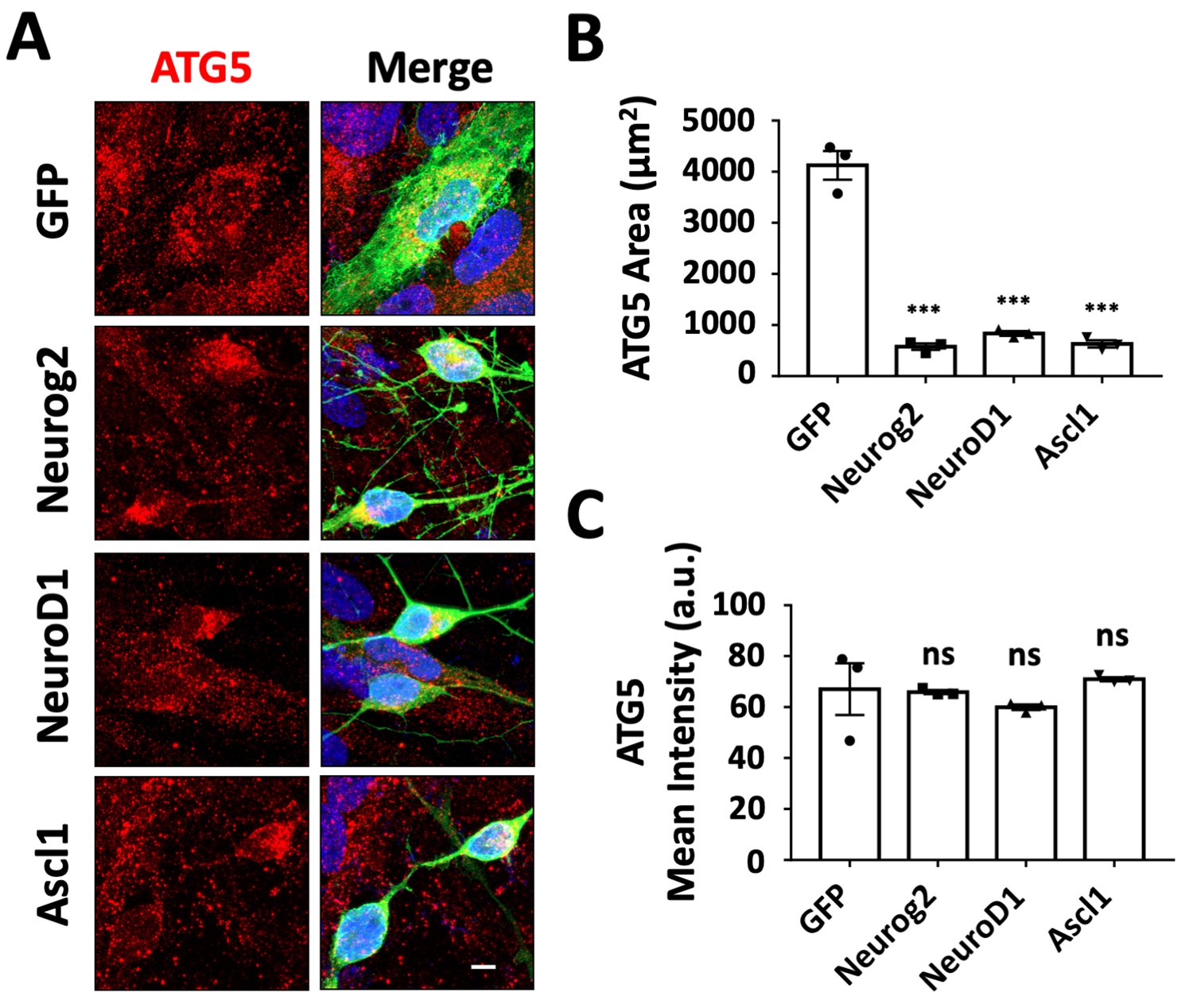

**Figure S5. Examination of autophagy/lysosomes during neuronal conversion of human GBM cells**

**A,** Representative images illustrating the distribution and morphological changes of autophagy/lysosomes (ATG5, red) during neuronal conversion of U251 human glioblastoma cells. Scale bars, 10 μm.

**B-C,** Quantification analyses of ATG5 covered area (**B**) and mean intensity (**C**) in transduced U251 cells at 30 dpi. n ≥ 150 cells from triplicate cultures. Data were represented as mean ± SEM and analyzed by one-way ANOVA followed with Dunnett’s test. *** p < 0.001.

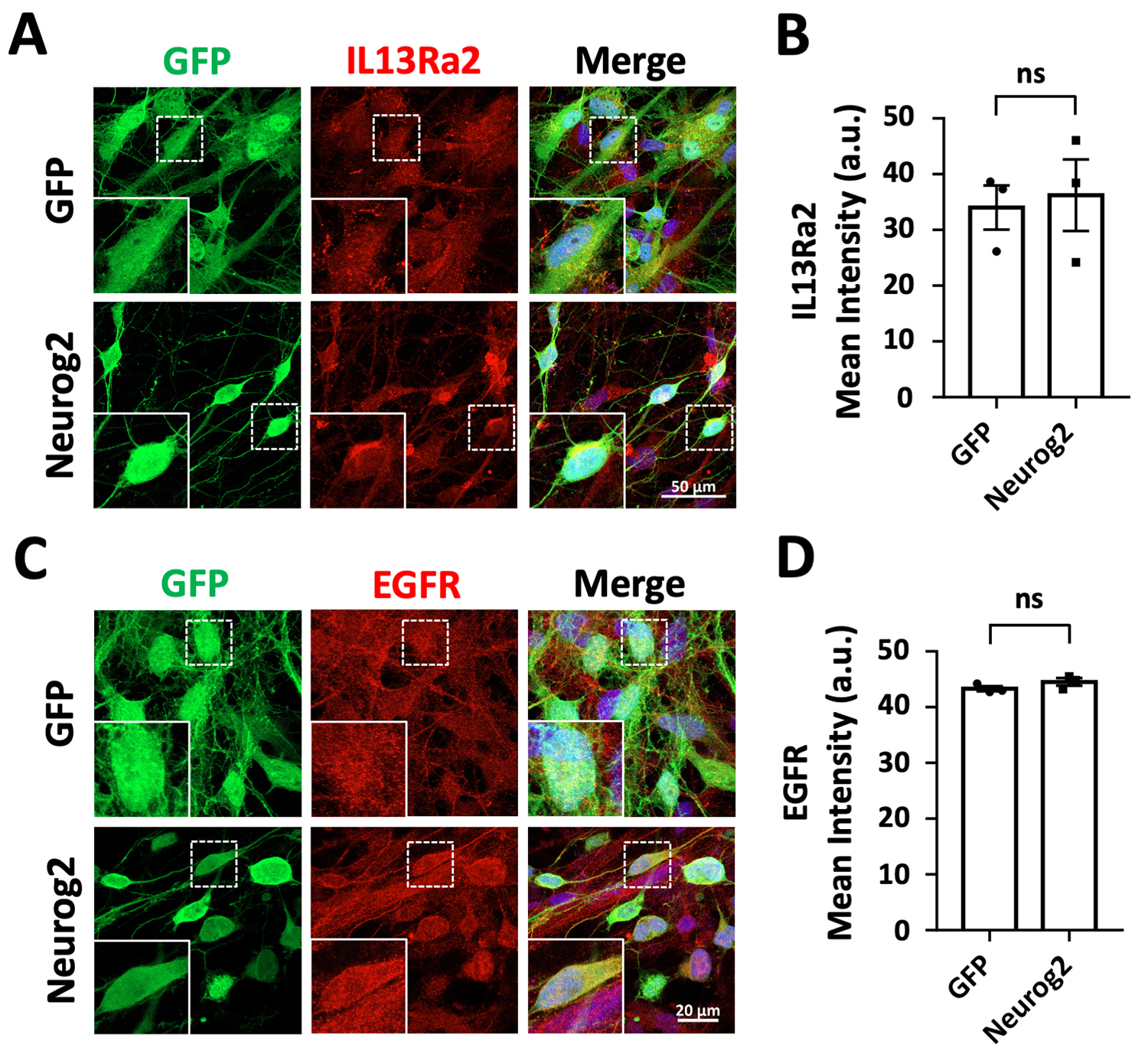

**Figure S6. Investigation of cancer makers in Neurog2-converted neurons from human glioblastoma cells**

**A,** Representative images showing the immunostaining of glioma restricted receptor IL13Ra2 (red) in U251 human glioblastoma cells expressing Neurog2 or GFP (green) at 20 dpi. Scale bars, 50 μm.

**B,** Quantification analyses of IL13Ra2 mean intensity in transduced U251 cells at 20 dpi.

**C,** Immunostaining of general cancer marker EGFR (red) in U251 human glioblastoma cells expressing Neurog2 or GFP (green) at 20 dpi. Scale bars, 50 μm.

**D,** Quantification analyses of EGFR mean intensity in transduced U251 cells at 20 dpi.

Data were represented as mean ± SEM and were analyzed by Student’s *t*-test. n ≥ 40 cells from triplicate cultures.

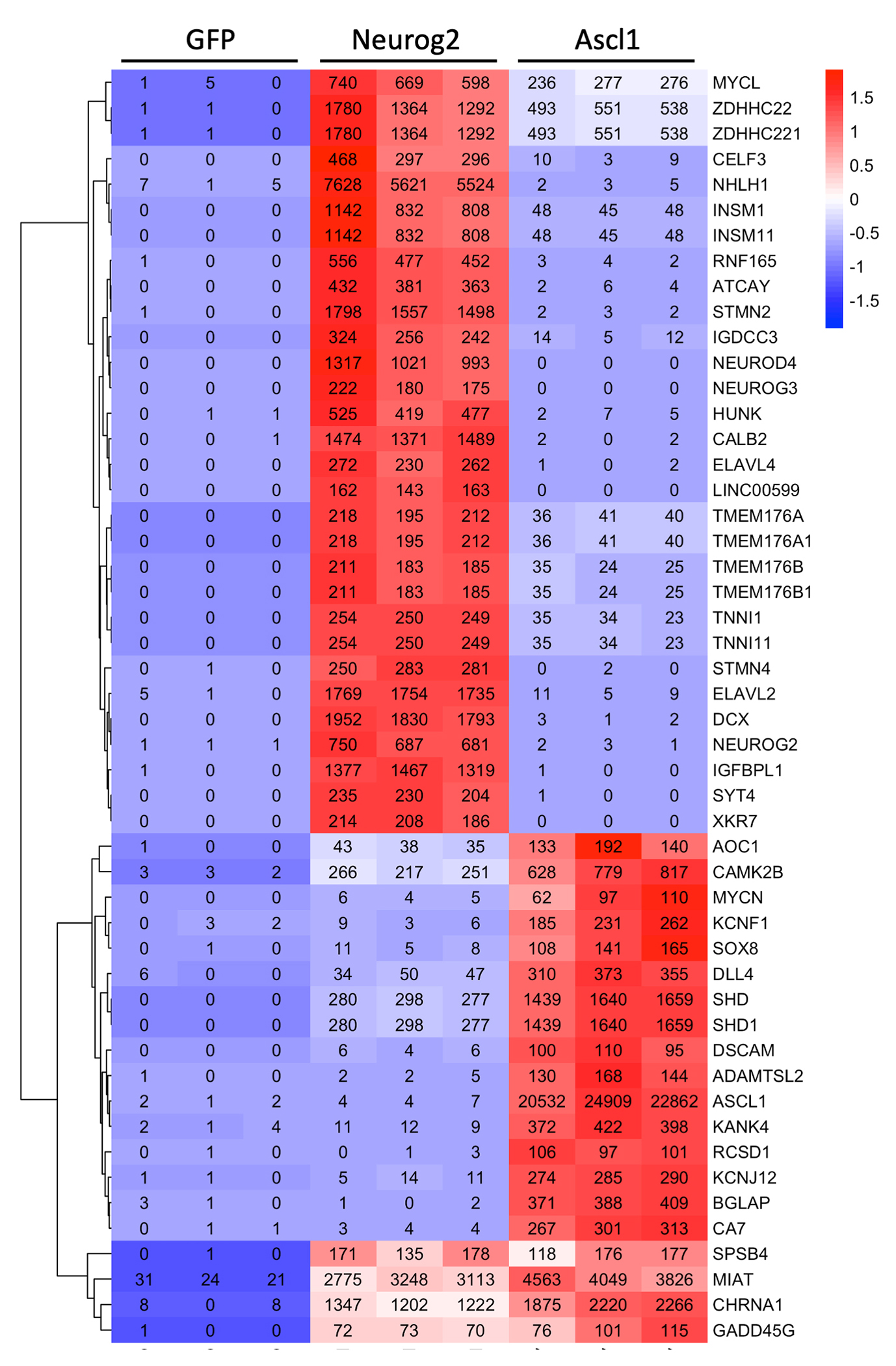

**Figure S7. Top upregulated DEGs in response to Neurog2 or Ascl1 overexpression in human glioblastoma cells**

A heat map with hierarchical clustering showing the top 25 upregulated DEGs (sorted by foldchange, > 100 normalized read counts in at least one sample) in response to Ascl1 or Neurog2 overexpression in U251 human glioblastoma cells. Color scaled within each row. Normalized read count values were presented.

**Supplemental Materials and Methods**

**Plasmid Construction and Retrovirus Production**

The mouse NeuroD1 plasmid was constructed from our PCR product according to a template of the pAd NeuroD-I-nGFP (Zhou et al., 2008) (Addgene) and inserted into a pCAG-GFP-IRES-GFP retroviral vector (Zhao et al., 2006) (gift of Dr. Fred Gage) to generate pCAG-NeuroD1-IRES-GFP. Viral particles were packaged in gpg helper free human embryonic kidney (HEK) cells to generate vesicular stomatitis virus glycoprotein (VSV-G)-pseudo typed retroviruses encoding neurogenic factors in CellMax hollow fiber cell culture system (Spectrum Laboratories). The titer of GFP, Neurog2-GFP, NeuroD1-GFP, Ascl1-GFP viral particles in virus-containing medium was about 1 x 10^3^ pfu/ml, 1 x 10^5^ pfu/ml, 1 x 10^4^ pfu/ml, 9 x 10^4^ pfu/ml respectively. The titer of GFP, Neurog2-GFP viral particles in concentrated viruses was about 2 x 10^5^ pfu/ml, 1 x 10^7^ pfu/ml respectively. Viral titer was determined by HEK cell transduction. For *in vitro* infection, around 500 ul GFP, 5 ul Neurog2-GFP, 50 ul NeuroD1-GFP, 5 ul Ascl1-GFP virus-containing medium was added to each well (24-well plates) to achieve comparable titer. For *in vivo* experiments, 2 ul GFP or 2 ul Neurog2-GFP (50 x diluted) concentrated retroviruses were mixed with human glioblastoma cells respectively for transplantation.

**Immunocytochemistry**

Cultured cells were fixed in 4% Paraformaldehyde (PFA) in PBS for 15 min at room temperature. Cells were washed three times by PBS and incubated in blocking buffer (5% normal donkey serum, 0.05% Triton X-100 in PBS) for 40min. Cells were incubated with primary antibodies in blocking buffer overnight at 4°C. The next day, cells were washed three times by 0.05% Triton X-100 in PBS and incubated with appropriate secondary antibodies conjugated to Alexa Fluor 488, Alexa Fluor 546, Alexa Fluor 647 (1:1000, Molecular Probes) for 1hr at room temperature. After three times washing with 0.05% Triton X-100 in PBS, coverslips were mounted onto a microscope slide (75 x 25 x 1 mm, VWR) with a mounting solution containing DAPI (Invitrogen). Slides were first examined with a revolve microscope (Echo, Revolve R4) and further analyzed with a confocal microscope (Zeiss LSM 800). Images were acquired and analyzed using ZEN software (Zeiss).

**RNA isolation, reverse transcription and RT-PCR**

RNA isolation from cultured cells were performed at desired time points using a NucleoSpin® RNA kit (Macherey-Nagel) following the manufacturer’s protocols. Reverse transcription was performed using 5 x qScriptTM cDNA SuperMix (Quanta Biosciences) from isolated RNA samples. PerfeCTaTM SYBR® Green SuperMix, ROXTM (Quanta Biosciences) was used for RT-PCR. GAPDH was used as the internal control. Each sample has three replicates for each target. The sequences of all primers were listed in Table 2.

**Western blotting**

Cells were lysed, fractionated by 10% SDS-Tris glycine and transferred to a 45 µm PVDF membranes. The membranes were blocked and incubated with the primary antibodies against GSK3β (27C10) (Rabbit, 1:1000, Cell Signaling 9315), and GAPDH (Rabbit, 1:5000, Sigma G9545) at 4 °C overnight. After washing, it was incubated with secondary antibodies (1:15000, Gt anti Rb 800, P/N 925-32210). Scanning was performed using LiCOR Odyssey Clx.

**Patch-Clamp Recordings in Cultured Cells**

For the converted neurons, whole-cell recordings were performed using Multiclamp 700A patch-clamp amplifier (Molecular Devices, Palo Alto, CA) as described before (Deng et al., 2007), and the chamber was constantly perfused with a bath solution consisting of 128 mM NaCl, 30 mM glucose, 25 mM HEPES, 5 mM KCl, 2 mM CaCl2, and 1 mM MgCl2. The pH of bath solution was adjusted to 7.3 with NaOH, and osmolarity was at 315–325 mOsm/l. Patch pipettes were pulled from borosilicate glass (3–5 MΩ) and filled with a pipette solution consisting of 135 mM KCl, 5 mM Na-phosphocreatine, 10 mM HEPES, 2 mM EGTA, 4 mM MgATP, and 0.5 mM Na2GTP (pH 7.3, adjusted with KOH). The series resistance was typically 10–30 MΩ. For voltage-clamp experiments, the membrane potential was typically held at −70 or −80 mV. Data were acquired using pClamp 9 software (Molecular Devices, Palo Alto, CA), sampled at 10 kHz, and filtered at 1 kHz. Na+ and K+ currents and action potentials were analyzed using pClamp 9 Clampfit software. Spontaneous synaptic events were analyzed using MiniAnalysis software (Synaptosoft, Decator, GA). All experiments were conducted at room temperature.

**Mitochondrial tracker incubation**

MitoTracker™ Red CMXRos (Invitrogen) was used to show mitochondrial morphology and distribution. MitoTracker was diluted with culture medium to a final concentration of 500 nM. Cells were incubated with MitoTracker for 1hr and then fixed with 4% PFA. This was followed by regular immunohistochemistry protocol.

**BrdU labeling and cell proliferation assays**

Cell proliferation was examined by BrdU incorporation. BrdU was added in cell culture medium (10 mM) as indicated durations. In desired time points, cells were fixed in 4% PFA for 15 minutes, and then treated with 2M HCl for 1 hour in room temperature, washed by PBS for 3 times with 5 min each time. This was followed by blocking and sequential incubations with anti-BrdU antibody (rat, 1:1000, Accurate Chemical, Westbury, NY, USA) and corresponding secondary antibody.

**Data and statistical analysis**

Cell counting and the fluorescence intensity were performed in a single blind way with randomly chosen fields of random chosen pictures and analyzed by Image J software. Data are represented as mean ± SEM. Multiple group comparisons were performed with two-way ANOVA followed with Dunnett’s test. Two group comparisons were performed with Student’s *t* test.

| Table 1. Antibodies used for Immunostaining | | | | |
| --- | --- | --- | --- | --- |
| Antibodies | **Species** | **Dilution** | **Company** | **Catalog No.** |
| Polyclonal anti-NEUN | Guinea pig | 1:1000 | Millipore | ABN90P |
| Polyclonal anti-MAP2 | Rabbit | 1:2000 | Millipore | AB5622 |
| Polyclonal anti-MAP2 | Chicken | 1:2000 | Abcam | AB5392 |
| Monoclonal anti-Tuj1 | Mouse | 1:1000 | COVANCE | MMS-435P |
| Polyclonal anti-Tbr1 | Rabbit | 1:600 | Abcam | AB31940 |
| Polyclonal anti-Prox1 | Rabbit | 1:1000 | ReliaTech GmbH | 102-PA32 |
| Polyclonal anti-FoxG1 | Goat | 1:600 | Abcam | AB3394 |
| Monoclonal anti-Ctip2 | Rat | 1:600 | Abcam | AB18465 |
| Polyclonal anti-VGluT1 | Rabbit | 1:1000 | Synaptic Systems | 135302 |
| Polyclonal anti-SV2 | Mouse | 1:1000 | DSHB | SV2 |
| Polyclonal anti-GFP | Chicken | 1:1000 | Abcam | AB13970 |
| Polyclonal anti-GFAP | Chicken | 1:1000 | Millipore | AB5541 |
| Polyclonal anti-GFAP | Rabbit | 1:1000 | Millipore | AB5804 |
| Monoclonal anti-  Human Nuclei (HuNu) | Mouse | 1:1000 | Millipore | MAB1281 |
| Monoclonal anti-S100β | Mouse | 1:1000 | Abcam | AB66028 |
| Polyclonal anti-DCX | Goat | 1:500 | Santa Cruz | SC-8066 |
| Polyclonal anti-SOX2 | Rabbit | 1:1000 | Millipore | AB5603 |
| Polyclonal anti-Ki67 | Rabbit | 1:1000 | Abcam | AB15580 |
| Monoclonal anti-BrdU | Rat | 1:1000 | Accurate | OBT0030 |
| Polyclonal anti-GABA | Rabbit | 1:1000 | Sigma | A2052 |
| Monoclonal anti-GM130 | Mouse | 1:800 | BD | 610822 |
| Polyclonal anti-ATG5 | Rabbit | 1:600 | Novus | NB110-53818 |
| Polyclonal anti-EGFR | Rabbit | 1:600 | Santa Cruz | SC-1005 |
| Monoclonal anti-Nestin | Mouse | 1:800 | Neuromics | MO15012 |
| Polyclonal anti-IL13Ra2 | Goat | 1:600 | R&D | AF146 |
| Polyclonal anti-Olig2 | Rabbit | 1:600 | Millipore | AB9610 |
| Polyclonal anti-GAP43 | Rabbit | 1:800 | Abcam | AB16053 |
| Polyclonal anti-Connexin 43 | Rabbit | 1:800 | Abcam | AB11370 |
| Monoclonal anti-GSK3β | Rabbit | 1:800 | Cell Signaling | 9315 |
| Polyclonal anti-Vimentin | Rat | 1:1000 | R&D | MAB2105 |
| Monoclonal anti-NeuroD1 | Mouse | 1:1000 | Abcam | AB60704 |
| Polyclonal anti-Neurog2 | Rabbit | 1:600 | Abcam | AB154293 |
| Polyclonal anti-Ascl1 | Rabbit | 1:800 | Abcam | AB74065 |

| Table 2. Sequences of primers used in RT-PCR | |
| --- | --- |
| Primers | **Sequence** |
| DCX-F | TGCTTGGGCCTCACACTAGC |
| DCX-R | CATATACCGCAATCAAGGAAATACTC |
| Ascl1-F | GTCAAGTTGGTCAACCTGGG |
| Ascl1-R | CTCATCTTCTTGTTGGCCGC |
| NEUROD2-F | TCAGACATGGACTATTGGCAG |
| NEUROD2-R | GGGACAGGAAAGGGAACC |
| NEUROG2-F | ATTTGCAATGGCTGGCATCT |
| NEUROG2-R | CACAGCCTGCAGACAGCAAT |
| NEUROD1-F | CCTGCAACTCAATCCTCGGA |
| NEUROD1-R | GGCATGTCCTGGTTCTGCTC |
| GAPDH-F | TGGGCTACACTGAGCACCAG |
| GAPDH-R | GGGTGTCGCTGTTGAAGTCA |
| ASCL1-F | CAACGACTTGAACTCCATGGC |
| ASCL1-R | TTGGTGAAGTCGAGAAGCTCC |
| DLX2-F | CAACAACGAGCCTGAGAAGGAG |
| DLX2-R | GGAAACTGGAGTAGATGGTGCG |
